# High-fat diet in early life triggers both reversible and persistent epigenetic changes in the medaka fish (*Oryzias latipes*)

**DOI:** 10.1101/2023.04.27.538519

**Authors:** Yusuke Inoue, Yuta Suzuki, Yoshimi Kunishima, Terumi Washio, Shinichi Morishita, Hiroyuki Takeda

## Abstract

The nutritional status during early life can have enduring effects on an animal’s metabolism, although the mechanisms underlying these long-term effects are still unclear. Epigenetic modifications are considered a prime candidate mechanism for encoding early-life nutritional memories during this critical developmental period. However, the extent to which these epigenetic changes occur and persist over time remains uncertain, in part due to challenges associated with directly stimulating the fetus with specific nutrients in viviparous mammalian systems. In this study, we used medaka as an oviparous vertebrate model to establish an early-life high-fat diet (HFD) model. Larvae were fed with HFD from the hatching stages (one week after fertilization) for six weeks, followed by normal chow (NC) for eight weeks until the adult stage. We examined the changes in the transcriptomic and epigenetic state of the liver over this period. We found that HFD induces fatty liver phenotypes, accompanied by drastic changes in the hepatic transcriptome, chromatin accessibility, and histone modifications, especially in metabolic genes. These changes were largely reversed after the long-term NC, demonstrating the high plasticity of the epigenetic state in hepatocytes. However, we found a certain number of genomic loci showing non-reversible epigenetic changes, especially around genes related to cell signaling, liver fibrosis, and hepatocellular carcinoma, implying persistent changes in the cellular state of the liver triggered by early-life HFD feeding. Our data provide novel insights into the epigenetic mechanism of nutritional programming and a comprehensive atlas of the long-term epigenetic state in an early-life HFD model of non-mammalian vertebrates.

## Introduction

Early-life nutrition can have life-long effects on the metabolism of animals. This phenomenon is referred to as “nutritional programming” (Langley-Evans, 2009). It is generally accepted that animals have the ability to adapt to the energy-imbalanced environments imposed during early life and continually shift their metabolic states throughout their lives (Pfennig et al., 2010). Accordingly, if early nutritional programming does not correctly match the postnatal environment, it may cause metabolic disorders such as obesity, type 2 diabetes, and cardiovascular diseases (Barker, 2007; Langley-Evans, 2009). Liver serves as the central organ regulating the metabolic homeostasis of animals across feeding-fasting cycles (Bideyan et al., 2021; Lempradl et al., 2015), adapting its phenotype in response to metabolic states, such as the development of fatty liver (Brunt et al., 2015). Regarding nutritional programming, cohort studies and experimental studies using mammalian models have shown that maternal high-fat diet (HFD) feeding, obesity, and diabetes during gestation and suckling period are associated with an increased risk of non-alcoholic fatty liver disease (NAFLD) in the offspring (de Jesus et al., 2020; Gregorio et al., 2010; Huang et al., 2017; Kruse et al., 2013; M. Li et al., 2015; Oben et al., 2010; Thompson, 2020). Epigenetic modifications are one of the prime candidate mechanisms implicated in long-term nutritional programming of the liver (Costello and Schones, 2018), since they can be maintained across cell divisions over a long period of time (Moazed, 2011). Indeed, in some types of cells such as innate immunity cells (Netea et al., 2016), epidermal stem cells (Larsen et al., 2021; Naik et al., 2017), and mammary epithelial cells (dos Santos et al., 2015), transient external stimuli caused long-term changes in chromatin state even after external stimuli were removed. These persistent epigenetic changes are associated with enhanced transcriptional responses to recurrent stresses at later stages (Larsen et al., 2021; Naik et al., 2017; Netea et al., 2016). Many studies that examined the effects of maternal nutritional stimuli on livers of offspring have shown long-term changes in epigenetic modifications, especially in DNA methylation (Chen et al., 2022; de Jesus et al., 2020; Maude et al., 2021; Suter et al., 2014). However, in the case of mammalian models, it can sometimes be difficult to determine a causal relationship between maternal nutrition and phenotypes of the offspring, as there are multiple pathways through which an imbalance of maternal nutrition can affect the fetus, such as maternal-fetal nutritional transmission via placenta or milk, transmission of maternal inflammatory cytokines and hormones, placental inflammation, microbiota, and maternal nursing behaviors (M. Li et al., 2015; Wesolowski et al., 2017). Previous studies with adult mice tested whether HFD-induced changes in chromatin state persisted when fatty liver was resolved by long-term feeding with normal chow (NC) (Leung et al., 2016; Siersbæk et al., 2017) or bariatric surgery (Ahrens et al., 2013; Du et al., 2017). However, the results are not consistent across studies, possibly due to the differences in experimental conditions such as age, strains, nutritional compositions of diet, and schedules of HFD feeding.

So far, studies on nutritional programming have been conducted mainly with mammalian models including humans, and knowledge in other non-mammalian vertebrates has been very limited. Medaka (Japanese killifish, *Oryzias latipes*) was recently utilized as a vertebrate model to analyze the effects of nutrition (Asaoka et al., 2014). Indeed, HFD feeding in adult medaka was shown to reproducibly lead to the development of fatty liver, obese phenotypes characterized by enlarged adipose tissues, as well as elevation of serum glucose and triglyceride levels, similar to the results observed in mammals (Matsumoto et al., 2010). Importantly, medaka is oviparous; starts feeding soon after hatching (7 days post fertilization (dpf)), providing a promising platform with which to apply direct nutritional stimulation from early stages. In addition, medaka has several advantages such as high fecundity and availability of high-quality genome data, making it possible to track epigenetic states throughout life (Ichikawa et al., 2017; Iwamatsu, 2004; Kasahara et al., 2007; Takeda and Shimada, 2010). Furthermore, given the long evolutionary distance between teleosts and mammals, studies with medaka could provide insights into conserved and divergent mechanisms of nutritional adaptation among vertebrates.

As a first step toward understanding the effect of early-life nutritional environment on non-mammalian vertebrates, we examined medaka livers which had experienced HFD feeding at larval stages, and assessed which epigenetic changes were induced and whether they persisted even after shifting to NC. Specifically, we performed RNA-seq, ATAC (Assay for Transposase-Accessible Chromatin)-seq, and ChIP (Chromatin immunoprecipitation)-seq of histone modifications (H3K27ac, H3K27me3, and H3K9me3). Under our experimental conditions, medaka responds well to HFD histologically, transcriptionally, and epigenetically to develop fatty liver. While HFD-induced transcriptomic and epigenetic changes almost entirely returned to normal levels as fatty liver was reversed, we found that some changes persisted at several specific loci. We discuss these non-reversible gene loci in light of nutritional programming.

## Results

### Fatty liver was induced by early-life HFD feeding in medaka, and was reversed following a switch to NC

To investigate the effects of early-life HFD feeding, we reared medaka fish in the following conditions (Figure 1A). First, male fish were divided into two groups and were fed either NC or HFD for six weeks from the hatching stage (seven weeks of age, termed NC and HFD fish, respectively). Both groups of fish were then fed NC for eight weeks, the period during which it was expected that the fatty liver condition would reverse (15 weeks of age, termed NC-NC and HFD-NC fish, respectively). To prevent competition between individuals and thus to reduce individual fluctuations as much as possible, we reared fish alone (one fish per one glass or plastic tank) for the duration of the study (Figure 1B). We found that body weight and liver weight were increased in HFD fish, but the difference in both parameters became smaller between NC-NC and HFD-NC fish (Figure 1C). We dissected livers from the fish in each group and performed histological analyses to check liver phenotypes. We confirmed that HFD fish showed the typical phenotypes of fatty liver, characterized by lipid droplets visualized by HE staining and neutral lipid accumulation detected by Oil Red O staining (Figure 1D). However, we did not observe any inflammatory cell aggregates, hepatocyte ballooning, or fibrosis, suggesting that six weeks of HFD feeding induces simple steatosis. After eight weeks of NC, those phenotypes disappeared, and no clear histological differences were detected between NC-NC and HFD-NC livers (Figure 1D). Electron microscopy observation confirmed the accumulation of lipid droplets in the livers of HFD fish, and their disappearance after 8-weeks of NC (Figure 1E). Thus, we concluded that early-life HFD feeding effectively induced weight gain and fatty liver, which could both be reversed by subsequent long-term NC feeding in medaka.

**Figure 1:**
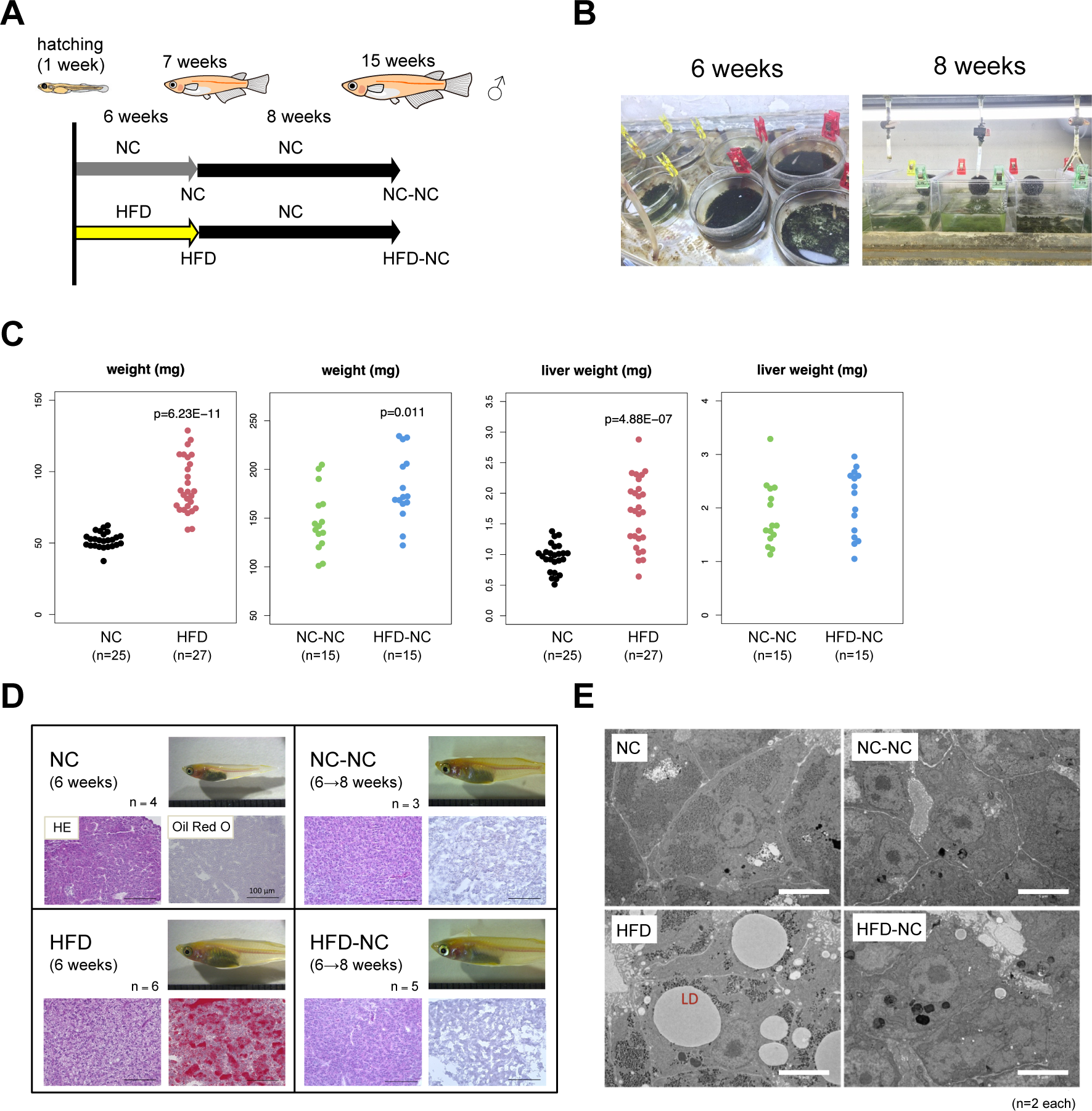
Early-life HFD feeding induces fatty liver in medaka, which is reversed by subsequent NC feeding. **(A)** Experimental design of early-life HFD feeding in medaka, where male fish were fed either HFD or NC for six weeks from hatching stage, followed by eight weeks of NC for both groups. **(B)** All fish were maintained individually to minimize variation of the speed of growth. **(C)** Body weight and liver weight at seven and 15 weeks of age. **(D)** Representative images of liver sections stained with H&E and Oil Red O, and **(E)** transmission electron microscopy (TEM) of liver sections from NC, HFD, NC-NC, and HFD-NC medaka. Scale bars: 100 µm in (D) and 5 µm in (E).

### Early-life HFD feeding caused drastic changes in hepatocyte gene expression, which were largely reversed by subsequent NC

We next investigated the effects of early and transient HFD on hepatic gene expression by RNA-seq. Since liver generally contains various types of cells other than hepatocytes (e.g., hepatic stellate cells, endothelial cells, cholangiocytes, erythrocytes, and immune cells), the analysis of whole-liver tissue would complicate the interpretation of results. Thus, we first established a transgenic medaka strain that expresses GFP specifically in hepatocytes. For this, we integrated the Tbait-hs-GFP cassette into the genomic region upstream of the transcriptional start site of *tdo2* (specifically expressed in hepatocytes) using the CRISPR-Cas9-mediated genome knockin method (Figure 2A) (Watakabe et al., 2018). Immunofluorescent staining confirmed that GFP was detected specifically in hepatocytes (Figure 2–figure supplement 1).

**Figure 2:**
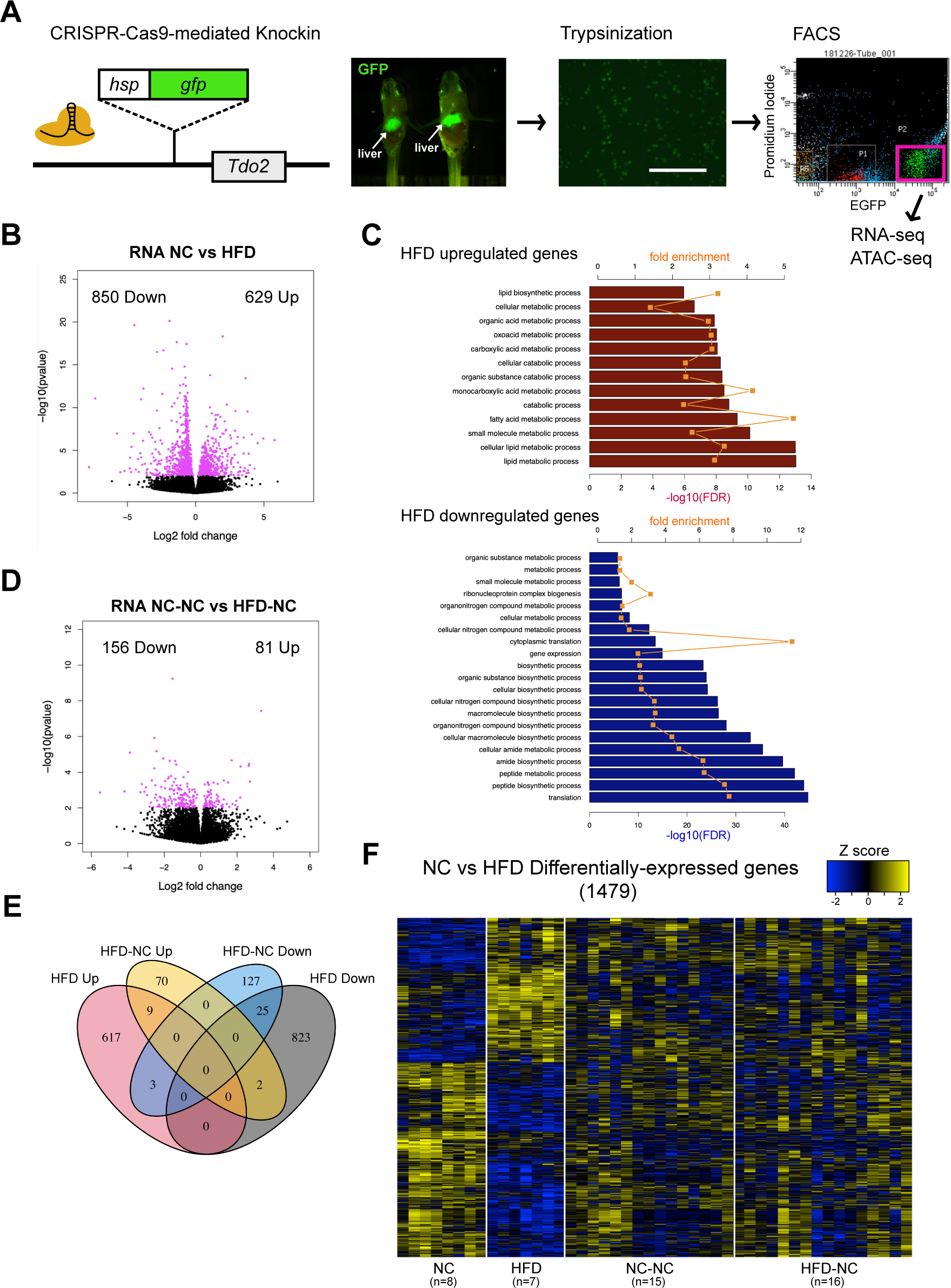
Early-life HFD feeding-induced changes in medaka hepatocyte gene expression are largely reversed by subsequent NC feeding. **(A)** A schematic of *tdo2*:GFP transgenic medaka construction using CRISPR-Cas9-mediated gene knockin and sorting of GFP-positive hepatocytes. Scale bar: 100 µm **(B)** RNA-seq results showing differentially expressed genes in hepatocytes of HFD fish compared to NC. X-axis: log2 fold change (HFD/NC) of expression values, y-axis: – log10 p-value. **(C)** Gene ontology analyses of upregulated (upper) and downregulated (lower) genes by HFD for biological processes using PANTHER 17.0. Fold enrichment (orange) and –log10 FDR (bar) are displayed. **(D)** Differentially expressed genes in hepatocytes of HFD-NC fish compared to NC-NC. **(E)** Venn diagram of differentially expressed genes at seven and 15 weeks of age. **(F)** A heatmap of differentially expressed genes at seven weeks of age. Log2-transformed, and Z-transformed, DESeq2 normalized read counts are displayed.

For transcriptome analysis, the transgenic fish were reared under the same conditions as shown in Figure 1A, then GFP-positive hepatocytes were isolated by fluorescence-activated cell sorting (FACS) and subjected to RNA-seq (Figure 2A, Figure 2–figure supplement 2). Each liver obtained from the fish in the four groups was individually analyzed (NC n=8; HFD n=7; NC-NC n=15; HFD-NC n=16). Despite individual rearing, we still observed large variation in gene expression between individuals, possibly caused by an accumulation of subtle differences during long-term rearing and/or inherent features of nutrition-related genes (Abu-Toamih Atamni et al., 2019). Thus, in the following analyses, we applied a less stringent threshold to identify differentially expressed genes. At seven weeks of age, 629 genes were up-regulated and 850 genes were down-regulated in HFD fish (Figure 2B, DESeq2, p-value < 0.01). Gene ontology analysis showed that upregulated genes in HFD hepatocytes were enriched for metabolic pathways, mainly lipid and fatty acid metabolic processes, while downregulated genes were enriched for translation and peptide biosynthetic processes (Figure 2C). Upregulation of lipid metabolism genes by HFD feeding was frequently reported in mammalian models of HFD feeding (Czech, 2017; Guan et al., 2018; Siersbæk et al., 2017; Soltis et al., 2017). It is unclear why translation-related genes (especially ribosomal protein genes) were downregulated, but it could be a result of endoplasmic reticulum (ER) stress inhibiting protein synthesis, which was also frequently reported in livers of HFD-fed mice (Lebeaupin et al., 2018). On the other hand, only 81 genes were up-regulated and 156 genes were down-regulated in HFD-NC hepatocytes, relative to NC-NC (Figure 2D, p-value < 0.01). When a more stringent threshold correcting for multiple testing at FDR < 0.1 was applied, only 15 genes were found to be differentially expressed between NC-NC and HFD-NC hepatocytes after 8 weeks of NC (Figure 2–figure supplement 3A). In addition, the number of genes persistently up-regulated or down-regulated following the eight weeks of NC is small (nine up-regulated and 25 down-regulated, Figure 2E, F). These data suggest that HFD feeding in early life stages induces drastic hepatic transcriptional changes similar to mammalian models, and that these changes are largely reversed by subsequent long-term NC feeding.

### Early-life HFD induced changes in the epigenetic state, which were largely reversed by subsequent NC

Previous studies have shown that some cell types, including innate immunity cells and epidermal stem cells, experience long-term epigenetic changes in response to transient inflammatory stresses, while transcriptional activity returns to near-normal levels (Larsen et al., 2021; Naik et al., 2017; Netea et al., 2016). To test if such persistent changes in the epigenetic state are induced by early-life HFD in medaka hepatocytes, we profiled genome-wide chromatin accessibility of FACS-isolated hepatocytes by ATAC-seq (Figure 2A). Each liver obtained from the fish in the four treatment groups was individually analyzed (NC n=7; HFD n=6; NC-NC n=9; HFD-NC n=6). We confirmed that the distribution of ATAC-seq signal exhibits expected patterns—enrichment at promoters and enhancers (Figure 3A). To identify ATAC-seq peaks that are differently regulated by diet, we initially called peaks for each dataset, merged the resulting peaks, and then selected reliable peaks based on their presence in at least six of the 28 samples. Then we counted the number of reads inside the defined peaks for each dataset and obtained differently accessible regions by DESeq2. Among the 30,484 open chromatin sites identified throughout the genome, 2,421 regions showed changes in chromatin accessibility soon after early-life HFD feeding (998 more accessible regions and 1,423 less accessible regions in HFD hepatocytes, DESeq2, p-value < 0.01, Figure 3B). Genomic annotation revealed that 70.7% of the differentially accessible peaks were located outside 2 kb on either side of the transcription start sites (TSS), which is a significantly higher proportion than the distribution observed for all peaks (55.4%, p-value < 2.2e-16, Fisher’s exact test). This suggests that there is a higher tendency for HFD-induced changes to affect chromatin accessibility of enhancers (Figure 3–figure supplement 1A). We next annotated all ATAC-seq peaks to their nearby genes (see Methods) and performed gene ontology analysis. Among the 30,484 peaks, 21,227 peaks were linked to at least one gene, and 13,622 genes were linked to at least one peak. Gene ontology analysis showed that genes related to monocarboxylic acid metabolic process were enriched in the set of genes with increased accessibility following HFD feeding (Figure 3C), consistent with the results of the RNA-seq analysis. Motif analyses using HOMER indicated that transcription factor motifs that regulate lipid metabolism (Nr2f6/Ppar, Srebp) were enriched, corroborating the observation that HFD feeding promotes lipid metabolic processes (Figure 3D). Genes associated with regulation of biological process, cell differentiation, and signaling were enriched among those with decreased accessibility in HFD hepatocytes (Figure 3C). Motifs of several liver-enriched transcription factors (e.g., forkhead box (Fox), GATA, HLF, C/EBP) and STAT were enriched for these peaks (Figure 3D). Eight weeks following the switch to NC, the number of differentially accessible peaks between dietary groups decreased, with 100 more accessible and 237 less accessible in HFD-NC compared to NC-NC (p-value < 0.01, Figure 3E, Figure 2–figure supplement 3B). Following the eight weeks NC, the number of peaks that remained persistently more-or less-accessible was limited (seven for up-regulated and 43 for down-regulated, Figure 3F), similar to the trend observed for transcriptional changes. Consequently, the chromatin state of metabolic genes, such as lipid metabolism genes, which exhibited altered chromatin accessibility after HFD feeding, was reversed by the subsequent NC (Figure 3G).

**Figure 3:**
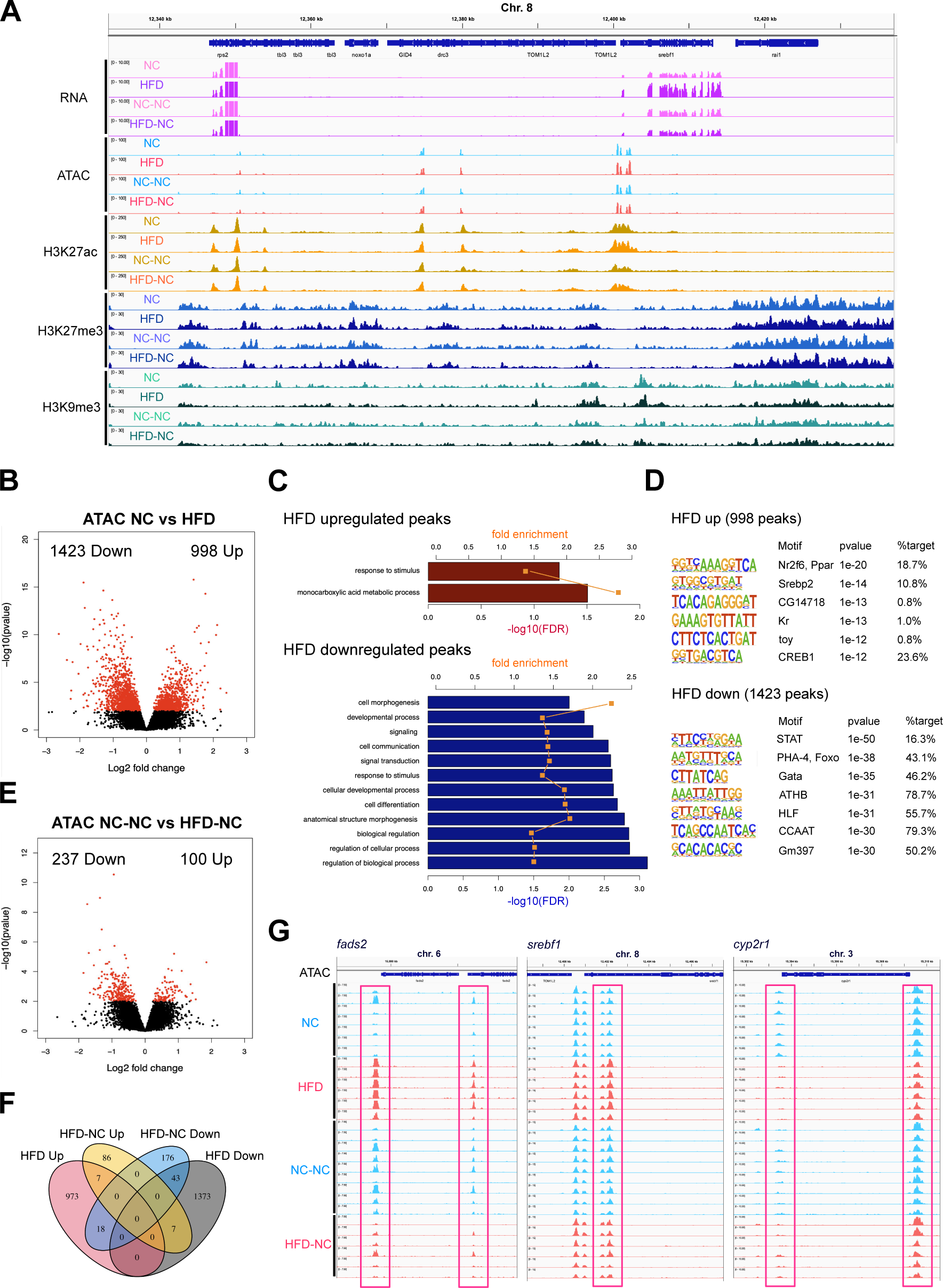
Early-life HFD feeding-induced changes in medaka hepatocyte chromatin accessibility are largely reversed by subsequent NC feeding. **(A)** A representative track view of RNA-seq, ATAC-seq, and ChIP-seq (H3K27ac, H3K27me3, and H3K9me3) signals for hepatocytes/livers for each of the dietary conditions. **(B)** Differentially accessible peaks in HFD fish hepatocytes compared to those in NC fish by ATAC-seq. X-axis: log2 fold change (HFD/NC) of normalized read counts within peaks, y-axis: –log10 p-value. **(C)** Gene ontology analysis of genes close to peaks with increased or decreased chromatin accessibility in HFD fish. Fold enrichment (orange) and –log10 FDR (bar) are displayed. **(D)** Motif analysis of peaks with increased (upper) and decreased (lower) accessibility, inferred by HOMER v4.11. **(E)** Differentially accessible peaks in HFD-NC fish hepatocytes compared to NC-NC fish. **(F)** Venn diagram of differentially accessible peaks at seven and 15 weeks of age. **(G)** Representative examples of ATAC-seq peaks at the promoters of *fads2*, *srebf1*, and *cyp2r1* genes.

We next performed ChIP-seq to profile the genome-wide distribution of three histone modifications (H3K27ac, H3K27me3, H3K9me3), which respectively mark active promoters and enhancers, repressive polycomb domains, and repressive heterochromatin. We first confirmed by western blot that the global levels of each histone modification were unchanged between dietary conditions (Figure 4–figure supplement 1). We also confirmed that the distribution of each histone modification exhibited expected patterns; H3K27ac was highly enriched in active gene promoters and enhancers and highly correlated with ATAC-seq signals, while H3K27me3 and H3K9me3 were enriched near repressed developmental genes and intergenic regions, and anti-correlated with ATAC-seq signals (Figure 3A, Figure 4–figure supplement 2A). To facilitate immunoprecipitation from a sufficient number of cells, three to seven livers were mixed together for each of the following ChIP-seq analyses, and three biological replicates were carried out.

**Figure 4:**
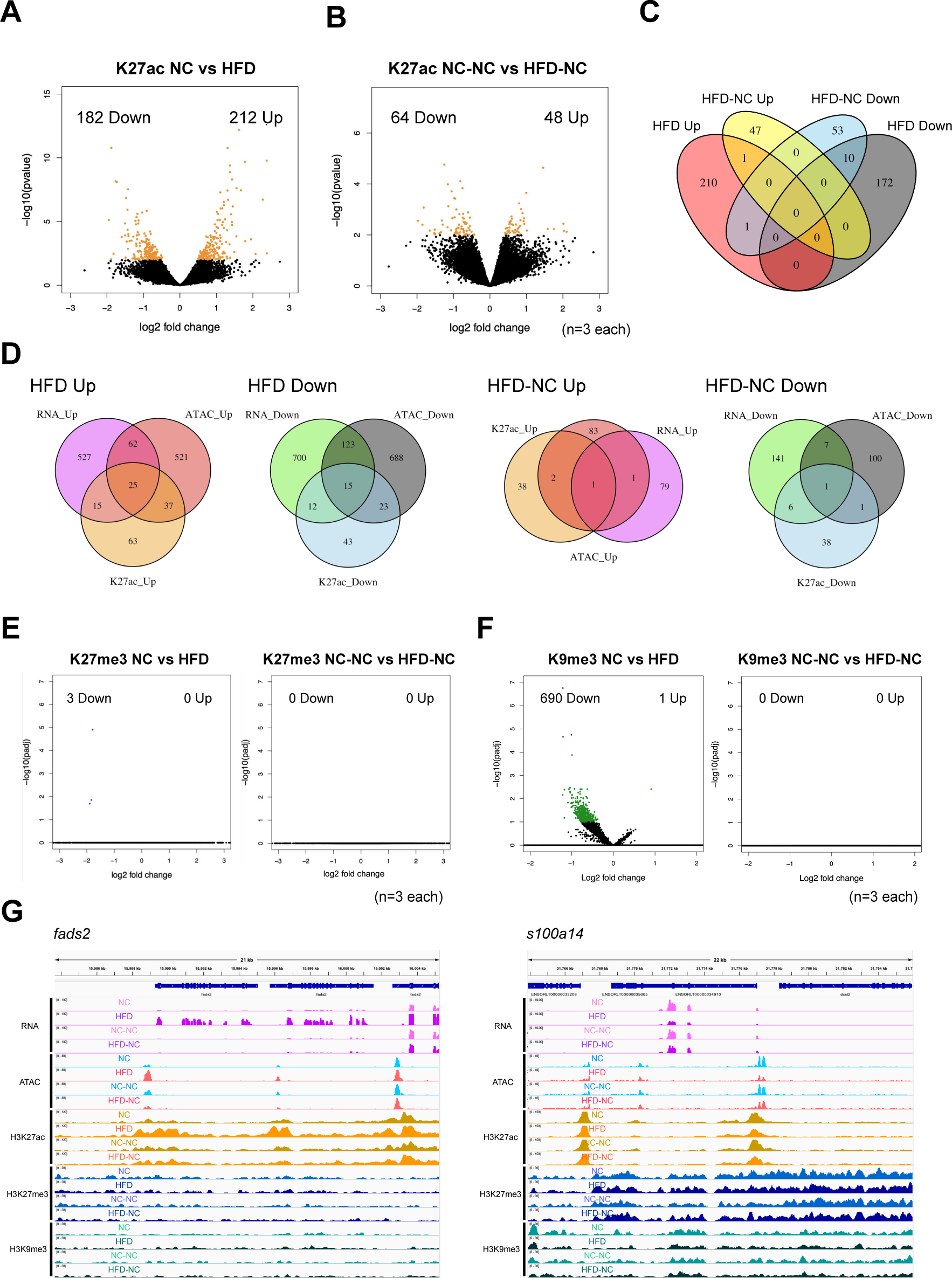
Early-life HFD feeding-induced changes in medaka liver histone modifications are largely reversed subsequent NC feeding. **(A)** Differentially-H3K27ac enriched peaks in livers of HFD-fed fish compared to NC by ChIP-seq. X-axis: log2 fold change (HFD/NC) of normalized read counts within peaks, y-axis: –log10 p-value. **(B)** Differentially-H3K27ac-enriched peaks in livers of HFD-NC fish compared to NC-NC. **(C)** Overlaps of differentially-H3K27ac-enriched peaks at seven and 15 weeks of age. **(D)** Overlaps of differentially expressed, differentially accessible, and differentially-H3K27ac-enriched genes, at seven and 15 weeks of age. **(E)** Differentially-H3K27me3-enriched 4-kb bins in livers of HFD fish relative to NC (left) and HFD-NC relative to NC-NC (right). **(F)** Differentially-H3K9me3-enriched 4-kb bins in livers of HFD fish relative to NC (left) and HFD-NC relative to NC-NC (right). **(G)** Representative examples of reversible peaks (i.e., genomic regions where epigenetic states were changed soon after HFD feeding but were returned to normal levels by the following NC feeding). Left, *fads2*; right, *s100a14*.

We identified 394 differentially-enriched peaks of H3K27ac soon after early-life HFD feeding (DESeq2, p-value < 0.01, Figure 4A, Figure 2–figure supplement 3C). Similar to the ATAC-seq results, differentially enriched peaks were predominantly located on enhancers (80.2% for differential peaks compared to 62.9% for all peaks, p-value = 3.64e-14, Fisher’s exact test, Figure 3–figure supplement 1B). Gene ontology analyses of nearby genes revealed that genes related to fatty acid metabolism were enriched for HFD-upregulated peaks, consistent with the results from the gene expression and chromatin accessibility analyses (Figure 4–figure supplement 3A). The number of differential peaks decreased to 112 after long-term reversal to NC (Figure 4B, Figure 2–figure supplement 3C), and only a limited number of persistent peaks were observed (one for up- and 10 for down-regulated in HFD-NC, Figure 4C), indicating that most changes in H3K27ac signal also returned to normal levels following the switch to NC (Figure 4–figure supplement 3B). Comparison of RNA-seq, ATAC-seq, and H3K27ac ChIP-seq results showed that changes in gene expression, chromatin accessibility, and H3K27ac signal were highly correlated with each other soon after HFD feeding, but the correlation weakened after long-term NC (Figure 4D, Figure 4–figure supplement 2B, Supplementary File 1).

Finally, we focused on the repressive histone modification marks of H3K27me3 and H3K9me3. However, due to their broad distribution pattern, we were unable to identify clear peaks. As an alternative, we divided the entire genome into 4-kb bins (resulting in 183,528 bins in total), counting the reads inside each bin and performing differential analysis using DESeq2. Surprisingly, we did not detect any regions differentially enriched for H3K27me3 soon after HFD feeding (Figure 4E), suggesting that the distribution pattern of H3K27me3 is not strongly influenced by HFD feeding. On the other hand, for H3K9me3, we identified 690 bins with significantly decreased ChIP read density in HFD livers (DESeq2, padj < 0.1, Figure 4F). Despite moderate enrichment for regulation of metabolic process in gene ontology analysis of nearby genes (Figure 4–figure supplement 4A), changes in H3K9me3 enrichment did not correspond to gene expression (Figure 4–figure supplement 4B) or correlate with chromatin accessibility (Figure 4–figure supplement 2B). Notably, after switching to NC, H3K9me3 signal mostly returned to control levels (Figure 4F, Figure 4–figure supplement 4C). Taken together, our results indicate that early-life HFD feeding causes drastic changes in chromatin accessibility and histone modifications, particularly H3K27ac and H3K9me3. The majority of these changes were reversed following a switch to NC, except for the particular loci described below. Representative examples are shown in Figure 4G.

### Persistently altered loci induced by early-life HFD

Although a recovery of fatty liver was observed, differences in the chromatin state were still found at 337 loci between NC-NC and HFD-NC fish at 15 weeks of age (Figure 3E). Gene ontology analysis revealed that these loci were enriched with genes related to cell surface receptor signaling pathway and cell differentiation (Figure 5A), suggesting potential changes at the levels of cellular or tissue properties in the livers that had experienced HFD, such as cell-cell interaction and differentiation. The 337 loci were a combination of persistently changed (i.e., difference in chromatin accessibility was observed at both seven and 15 weeks of age) and transiently changed (i.e., difference in chromatin accessibility was observed only at 15 weeks of age) ATAC-seq peaks. We focused on the 50 persistent peaks (Figure 3F, Figure 5B, Figure 5–figure supplement 1A). While differences at each locus may be sometimes subtle, we reason that some of these loci could truly maintain altered chromatin accessibility induced by early-life HFD feeding, if they are accompanied by consistent changes in gene expression and H3K27ac signal. In this context, at least 16 peaks among the 50 persistent peaks, were accompanied by consistent changes in gene expression and/or H3K27ac at seven and/or 15 weeks of age (Figure 5B, Figure 5–figure supplement 1A). These include peaks showing long-term changes in expression of nearby genes (*raftlin*, *hhla2b.1*, and *epha5*, Figure 5C), and peaks showing persistent epigenetic changes while gene expression returned to normal levels (*cdh18*, *gadd45a*, and two unannotated genes, Figure 5D). Intriguingly, we found that some of the 50 peaks are located close to genes related to cell signaling (e.g., *dscam1*, *epha5*, *ptprfb*, *hhla2b*, *cdh18*, *rasgef1bb*, and *raftlin*), and are associated with liver fibrosis and hepatocellular carcinoma (HCC) (e.g., *epha5*, *raftlin*, *hhla2b1*, and *gadd45a*, Figure 5B-D). Additionally, we identified 11 peaks showing persistent changes in H3K27ac signal, and 10 of them showed consistent changes in expression or ATAC-seq signal at either time point (Figure 4–figure supplement 3C, Figure 5–figure supplement 1B). Therefore, despite the nearly reversible chromatin state triggered by HFD feeding, there are a number of loci showing persistent changes in chromatin state following a switch to long-term NC. These are candidate genomic loci responsible for long-term epigenetic memory of early-life HFD feeding (see Discussion).

**Figure 5:**
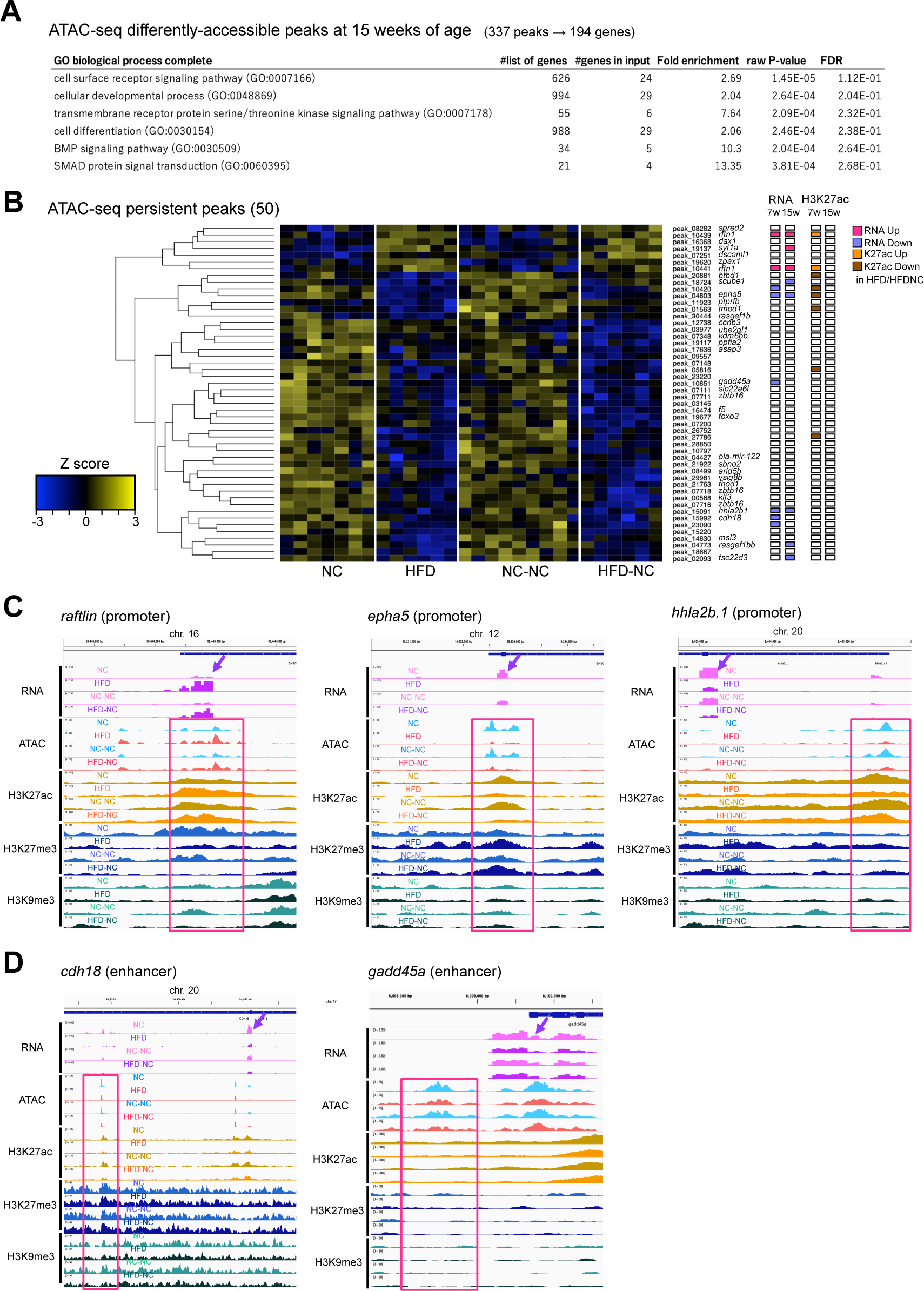
Genes showing persistent changes in epigenetic state following long-term NC. **(A)** Gene ontology analysis of genes close to differentially accessible ATAC-seq peaks between HFD-NC and NC-NC fish (194 genes from 337 peaks). **(B)** A heatmap of persistent ATAC-seq peaks after reversal to NC (50 peaks in total). Log2-transformed, and Z-transformed, DESeq2 normalized read counts of ATAC-seq at each peak are displayed. DESeq2 results of RNA-seq and H3K27ac ChIP-seq of nearby genes/peaks are displayed on the right (p-value < 0.01). **(C, D)** Representative examples of ATAC-seq peaks showing persistent changes in chromatin accessibility, accompanied by long-term changes in expression of nearby genes (C) or gene expression that returned to normal level after a switch to NC (D).

## Discussion

Several studies conducted in mammals have shown that overnutrition during development and growth periods increases susceptibility to obesity, diabetes, and fatty liver disease in later life (de Jesus et al., 2020; Gregorio et al., 2010; Huang et al., 2017; Kruse et al., 2013; M. Li et al., 2015; Oben et al., 2010; Thompson, 2020). Epigenetic modifications have been proposed as the mechanism underlying the long-term memory of early-life overnutrition (Costello and Schones, 2018). However, the extent to which epigenetic changes induced by early-life overnutrition persist and affect later metabolic programming phenotypes remains controversial across studies (Ahrens et al., 2013; Du et al., 2017; Kim et al., 2019; Leung et al., 2016; Siersbæk et al., 2017). To avoid the complexity of mammalian systems (M. Li et al., 2015; Thompson, 2020; Wesolowski et al., 2017), we developed a new system using medaka as an oviparous animal model, and in this study we tested whether HFD feeding during the early growth period can effectively cause fatty liver phenotypes and to what extent induced transcriptional and epigenomic changes persist. To our knowledge, this is the first study to thoroughly analyze the long-term effect of early-life nutritional environment on the epigenetic state in non-mammalian vertebrates.

Our data demonstrate that HFD induces drastic changes in the hepatic transcriptome, chromatin accessibility, and histone modifications especially in metabolic genes, and that subsequent long-term NC feeding returned most of the changes to normal levels. Our results suggest that the hepatic transcriptome and epigenome of medaka are highly adaptable to changes in the feeding environment. However, we also notably found genomic loci which showed persistent epigenetic changes triggered by early-life HFD feeding (discussed later).

Our results are consistent with several recent studies in adult mice investigating the persistence of HFD-induced epigenetic changes following weight loss by long-term NC or bariatric surgery (Du et al., 2017; Kim et al., 2019; Siersbæk et al., 2017). For example, Siersbæk et al. demonstrated that seven weeks of HFD feeding caused fatty liver accompanied by drastic changes in the hepatic transcriptome and H3K27ac at enhancers, which returned to levels comparable with control mice after five weeks of NC and recovery from fatty liver (Siersbæk et al., 2017). However, Siersbæk et al., found that chromatin accessibility was not affected by HFD feeding, based on DNase-seq analysis. On the other hand, our data based on ATAC-seq analysis demonstrates that HFD feeding causes drastic changes in chromatin accessibility. The reason for the discrepancy is unclear, but one possibility is the difference in the timing of HFD feeding (i.e., Siersbæk et al. fed HFD to 12 weeks old adult mice, while we started HFD feeding from the hatching stage in medaka). Similarly, Du et al. surveyed the persistence of HFD-induced changes in chromatin accessibility long after vertical sleeve gastrectomy by FAIRE-seq, and found that chromatin accessibility returned to normal levels after weight loss (Du et al., 2017). In addition, Kim et al. tested the persistence of HFD-induced changes in DNA methylation following nine weeks of NC, and again showed that the majority of hyper- and hypomethylated CpGs were reversed after weight loss, although ∼5.7% of CpGs (152 CpG in total) showed persistent changes, similar to our results (Kim et al., 2019). In contrast, Leung et al. demonstrated that HFD-induced changes in chromatin accessibility persisted eight weeks after a switch to NC; however, this difference can be explained by the incomplete reversal of obese phenotypes in their study (i.e., heavier body weight and more lipid accumulation in livers in the HFD-NC mouse group) (Leung et al., 2016). Taking into account these previous studies, it is reasonable to speculate that HFD-induced changes in epigenetic state are largely reversible upon recovery from fatty liver phenotypes. The high plasticity of epigenetic states in hepatocytes seems plausible, as hepatocytes need to dynamically regulate metabolic genes according to the feeding/fasting cycle, to maintain whole-body metabolic homeostasis (Bideyan et al., 2021). Indeed, dynamic circadian regulation of enhancer RNA production and H3K27ac accumulation was reported in livers of mice (Fang et al., 2014).

So far, studies of nutritional programming have been mainly conducted in mammals, because of clinical relevance. However, the long-term effects of early-life nutritional intervention have also been widely investigated in teleosts, with the aim of producing high-quality farmed fish at low cost (Panserat et al., 2019). The strategy is based on the concept that the first feeding stage, the transition from endogenous-to exogenous feeding, exhibits a high degree of molecular plasticity for intermediary metabolism, leading long-term metabolic traits that persist throughout life. One example is feeding of an early-life carbohydrate-rich (CH-rich) diet to carnivorous fish (Fang et al., 2014; Geurden et al., 2014, 2007; Gong et al., 2015; Rocha et al., 2016), which generally exhibit poor utilization of carbohydrates due to their weak glucose metabolism such as glucose intolerance, poor carbohydrate digestion, absorption, and utilization (Kamalam et al., 2017). In these studies, a CH-rich diet at first feeding resulted in long-term changes in carbohydrate metabolism in response to a secondary CH-rich diet challenge, such as lower expression of gluconeogenesis genes (zebrafish (Fang et al., 2014) and Siberian sturgeon (Gong et al., 2015)), increased expression of amylase and maltase (rainbow trout (Geurden et al., 2007)), and improved uptake and catabolism of carbohydrates (gilthead seabream (Rocha et al., 2016)). Another example is a plant-based diet at the initial phase of feeding (Balasubramanian et al., 2016; Clarkson et al., 2017; Geurden et al., 2013; Michl et al., 2017; Vera et al., 2017), which is rich in various secondary metabolites and poor in long-chain poly-unsaturated fatty acids (PUFAs), leading to poor growth in some carnivorous and/or marine species (Panserat et al., 2019; Vera et al., 2017). A first feeding of a plant-based diet to rainbow trout and Atlantic salmon resulted in improved growth rate and energy utilization during the secondary plant-based diet challenge, compared to fish fed a marine based diet (Clarkson et al., 2017; Geurden et al., 2013). Microarray expression analysis in the liver of juvenile Atlantic salmon challenged with a plant-based diet revealed the upregulation of genes involved in intermediary metabolism (e.g., glucose metabolism, fatty acid biosynthesis, and elongation), protein metabolism, and antioxidant defense of cells, suggesting an improved utilization of a plant-based diet at the molecular level (Vera et al., 2017). These examples clearly demonstrate that long-term changes can occur in the metabolism of fish. This somewhat contrasts with our results in medaka liver where the effects of early-life HFD are mostly reversible. However, it is worth noting that dietary habits are quite diverse among fish and this is reflected in the manner of nutritional adaptation; narrow (e.g., carnivorous and/or marine) vs omnivorous feeding freshwater fish (Medaka). To understand the mechanisms of nutritional programming in fish, it will be intriguing to apply genome-wide epigenome profiling to the above-mentioned conditioned fish, and our study provides reference data for those future studies.

Although HFD-induced changes in chromatin state are largely reversible in medaka, we found a number of ATAC-seq peaks showing non-reversible changes long after a switch to NC (Figure 5, Figure 5–figure supplement 1). At least 16 peaks were accompanied by consistent changes in gene expression and/or H3K27ac at either seven and/or 15 weeks of age, suggesting that they underwent persistent epigenetic changes (Figure 5B, Figure 5–figure supplement 1A). Some peaks showed consistent changes in expression of nearby genes at both time points (e.g., *raftlin*, *hhla2b.1*, and *epha5*), suggesting that the epigenetic state of chromatin has a direct impact on genes near those peaks (Figure 5C). On the other hand, a majority of peaks were not associated with changes in gene expression at 15 weeks of age. We speculate that at least some of these peaks exhibit latent epigenetic changes, impacting nearby genes under some specific conditions. The fact that most of those persistent peaks were categorized as enhancers supports this idea (Figure 3–figure supplement 1), since chromatin accessibility at enhancers does not necessarily reflect gene expression (Klemm et al., 2019; Larsen et al., 2021; Netea et al., 2016). These peaks could be candidates with long-term epigenetic change triggered by early-life HFD feeding.

Interestingly, some of the persistently changed ATAC-seq peaks are located close to genes related to cell signaling and associated with liver fibrosis and hepatocellular carcinoma. For example, *epha5*, a persistently downregulated gene in terms of expression and chromatin accessibility in HFD-NC fish, has been reported to be associated with several types of cancers (S. Li et al., 2015), including hepatocellular carcinoma (Wang et al., 2019). Eph receptors, including Epha5, constitute the largest subfamily of receptor tyrosine kinases and mediate a variety of cellular processes such as cell migration, adhesion, proliferation, and differentiation, in response to their ligands ephrins (Kania and Klein, 2016; Shu et al., 2022). Ephrin/Eph signaling plays pivotal roles in various developmental contexts such as axon guidance, tissue boundary formation, and angiogenesis (Kania and Klein, 2016), but its dysregulation is frequently observed in various types of cancers (Shu et al., 2022). Similarly, *raftlin*, a persistent upregulated gene in this study, tends to be upregulated in patients of HCC compared to healthy controls (Sameh Boshra et al., 2019). Raftlin is one of the major lipid raft proteins and is known to function in adaptive immunity by mediating B-cell receptor (BCR) and T-cell receptor (TCR) signaling in lymphocytes, as well as in innate immunity by mediating uptake of ligand-bound Toll-like receptors (TLR) 3/4 via endocytosis in dendritic cells and macrophages (Matsumoto and Tatematsu, 2016; Saeki et al., 2003). Although the function of Raftlin in liver, as well as its association with HFD feeding and obesity, has not yet been tested, persistent upregulation of this gene could imply latent activation of innate immunity, which affects the propensity for liver inflammation and fibrosis. In addition, some other persistent genes have been implicated in liver damage and fibrosis (e.g., *dax1*, *gadd45a*, *tsc22d3*, and *miR-122*) (Flamini et al., 2021; Hsu et al., 2012; Tanaka et al., 2017; Yun et al., 2022) and HCC (e.g., *spred2*, *scube1*, *ptprfb*, *tmod1*, *asap3*, *zbtb16*, and *hhla2b1*) (Bera et al., 2014; Feng et al., 2020; Gao et al., 2023; Hui et al., 2015; Luo et al., 2021; Vi et al., 2008; Zhao et al., 2022). Thus, the persistently-affected genes identified in our study could represent latent changes in the cellular state of the liver that may contribute to the increased propensity for fibrosis and HCC development later in life as a result of early-life HFD feeding in medaka. Intriguingly, studies with mice also reported persistent changes in DNA methylation around genes involved in cell signaling and development in adult offspring of HFD-fed dams (Suter et al., 2014; Wankhade et al., 2017). They are associated with a higher propensity to develop liver fibrosis (Wankhade et al., 2017), suggesting that these may represent conserved features of long-term nutritional effects in mice and medaka. To reveal the roles of these persistent genes, further phenotypic analyses focusing on aged medaka, as well as analyses focusing on the interaction of hepatocytes with other types of liver cells (e.g., hepatic stellate cells, endothelial cells, and macrophages) are needed. Finally, further studies with non-mammalian vertebrates with various dietary habits will provide unique and conserved insights into the mechanism and evolution of nutritional adaptation.

## Methods

### Animals

Himedaka strain d-rR or *tdo2*:GFP transgenic medaka were used in this study. All experimental procedures and animal care were performed according to the animal ethics committee of the University of Tokyo. Fish eggs were collected and incubated at 28.5℃ for seven days until hatching. After then, fish were raised in a circular glass tank with 80 mm diameter and 50 mm depth individually (a single fish per a single tank) for six weeks to avoid competition between individuals. During this period, fish were fed either normal chow (Hikari rabo 130, Kyo-rin, approx. 20.0% calories from fat, 61.0% calories from protein and 19.0% calories from carbohydrate) or high-fat diet (High Fat Diet 32; CLEA Japan Inc., 56.7% calories from fat, 20.1% calories from protein and 23.2% calories from carbohydrate) twice a day. To prevent high mortality, a small amount of NC was supplemented to the HFD group fish for the morning feed. All glass cups were maintained in a large water bath, with a constant temperature of 29.0∼29.5℃. After seven weeks of age, fish were transferred to small plastic tanks with 130 mm width, 90 mm depth, and 85 mm height, individually (a single fish per single tank) in tap water and fed NC (Tetramin Super, Tetra, approx. 26.9% calories from fat, 58.7% calories from protein and 14.4% calories from carbohydrate) twice a day for eight weeks. Throughout fish rearing, room temperature was maintained at 27℃±1℃, with a strict light-dark cycle (lights turned on at 8:30 am and turned off at 10:00 pm). In all experiments, fish were sampled between 1:00 to 3:00 pm after undergoing a period of fasting that began in the afternoon of the preceding day.

### Histology

The fish were anesthetized with ice-cold water, and their livers were dissected for histological analyses. For hematoxylin and eosin (HE) staining, the liver tissue was fixed overnight at room temperature with gentle shaking using Bouin’s fixative. Afterward, the tissues were dehydrated with ethanol, cleared using Clear Plus (Falma), infiltrated with paraffin at 65℃, and embedded in it. 5 µm-thick sections were cut and stained with HE. For Oil Red O staining, the liver was fixed with 4% paraformaldehyde in PBS for overnight at 4℃ with gentle shaking. After washing twice with PBS, liver was transferred to 30% sucrose/PBS and rotated overnight at 4℃, and embedded in O.C.T compound (Sakura Finetek Japan). 10 µm-thick frozen sections were cut, dried, and stained with 0.18% Oil Red O/60% isopropanol for 10 min at 37℃.

### Transmission electron microscopy

The samples were fixed with 2% paraformaldehyde (PFA) and 2% glutaraldehyde in 0.1 M phosphate buffer (PB) pH 7.4 at 4℃ overnight. After washing with 0.1 M PB, liver tissues were postfixed with 2% osmium tetroxide (OsO_4_) in 0.1 M PB at 4℃ for 2h. After dehydration in graded ethanol solutions (50%, 70%, 90%, and 100%), livers were infiltrated with propylene oxide (PO), a 70:30 mixture of PO and resin, and embedded in 100% resin. Ultra-thin sections (70 nm-thick) were cut, mounted on copper grids, and stained first with 2% uranyl acetate, then with Lead stain solution (Sigma). The grids were observed by transmission electron microscopy (JEM-1400Plus; JEOL Ltd.) at an acceleration voltage of 100 kV. All procedures after fixation were performed by Tokai Electron Microscopy. Inc. (Aichi, Japan).

### Generation of *tdo2*:GFP transgenic medaka

*Tdo2*:GFP transgenic medaka was generated as described in Watakabe et al., 2018 (Watakabe et al., 2018). Donor plasmid (Tbait-hs-GFP) and a pDR274 plasmid containing sgRNA sequence targeting Tbait were kindly provided by Dr. Higashijima at the National Institute for Basic Biology (NIBB). Cas9 mRNA was synthesized by digesting the pCS2-Cas9 plasmid by NotI, followed by transcription using an mMESSAGE mMACHINE SP6 kit (Life Technologies). A sgRNA targeting *tdo2* was designed using CCTop (Stemmer et al., 2015), using the 1000 bp region upstream of the transcription start site of *tdo2* as a query. sgRNA was synthesized as follows; first, guide oligos were annealed by heating to 98℃ for 30 seconds, followed by ramping down to 25℃ at a rate of −0.1℃/sec. Then, 40 ng of pDR274e plasmid and ∼0.1 pmol of annealed oligos were mixed with Ligation High ver. 2 (TOYOBO) and BsaI-HF (NEB) and incubated at 37℃ for 5 min and 16℃ for 5 min for three cycles, to insert the annealed oligo to the pDR274e plasmid. After amplification of the template DNA by PCR, sgRNA was synthesized using NEB’s HiScribe T7 Quick High Yield RNA Synthesis Kit (NEB). For generation of transgenic fish, a cocktail of sgRNAs, donor plasmid, and Cas9 mRNA (9 ng/µl of sgRNA for digesting Tbait, 18 ng/µl of sgRNA for digesting genome DNA, 9 ng/µl of donor plasmid and 200 ng/µl of Cas9 mRNA) was injected into one-cell stage medaka embryos.

### Immunohistochemistry

Liver was dissected from *tdo2*:GFP medaka and fixed in 4% PFA/PBS for two hours at room temperature. After washing with PBS, the liver was dehydrated with ethanol, cleared using Clear Plus (Falma), infiltrated with paraffin at 65℃, and embedded in it. 5 µm-thick sections were cut and immunostained with anti-GFP antibody (living colors, Clontech, 632592) and secondary anti-rabbit IgG with Alexa fluor 488 (Thermo Fisher). Images were captured by Zeiss LSM700 confocal microscopy.

### Isolation of GFP-positive hepatocytes for RNA-seq and ATAC-seq

Livers were dissected from *tdo2*:GFP medaka and minced into small pieces on ice using a razor blade. Then, cells were dissociated into individual cells in 0.25% (w/v) trypsin (Nacalai Tesque)/PBS at 37℃ for 5 min, and dissociation was stopped by adding the same volume of 15% (v/v) FBS/Leiboviz’s L-15 (Life Technologies). After gentle pipetting 30 times and filtration using pluriStrainer Mini 40 µm (pluriSelect), cells were centrifuged again and resuspended with 15% FBS/L-15. GFP-positive cells (hepatocytes) were sorted using FACSAria III (BFD Biosciences). Dead cells were stained with propidium iodide (Life Technologies) and removed during cell sorting.

### RNA-seq

Total RNA was extracted from 50,000 of the sorted hepatocytes using RNeasy Mini Kit (Qiagen). mRNA was enriched by poly-A capture and mRNA-seq libraries were generated using KAPA Stranded mRNA-seq Kit (KAPA Biosystems). Libraries were sequenced using the Illumina HiSeq 1500 platform.

### ATAC-seq

ATAC-seq was performed as described in Buenrostro et al. 2015, with some modifications (Buenrostro et al., 2015). 50,000 of the sorted hepatocytes were washed with PBS and resuspended with ice-cold lysis buffer (10 mM Tris-HCl, pH 7.4; 10 mM NaCl; 3 mM MgCl_2_; 0.1% (v/v) Igepal CA-630) to extract nuclei. After centrifugation at 500×g, 4℃ for 10 min, nuclei pellet was resuspended with the transposition reaction mix (25 µl of 2x reaction buffer and 2.5 µl of Tn5 transposase from Nextera Sample Preparation Kit (Illumina), mixed with 22.5 µl nuclease-free water) and incubated at 37℃ for 30 min. Following transposition, DNA was purified using a Qiagen MinElute PCR Purification Kit (Qiagen) and eluted with 10 µl of Buffer EB. Two sequential PCRs were performed to enrich small DNA fragments. First, nine-cycles of PCR were performed using indexed primers from a Nextera Index kit (Illumina), and amplified DNA was size-selected to a size of <500 bp using AMPure XP beads (Beckman Coulter). Then, seven-cycles of PCR were performed using the same primers as the first PCR and purified by AMPure XP beads. Libraries were sequenced using the Illumina HiSeq 1500 platform.

### ChIP-seq

Liver chromatin lysate was prepared as described in Schmidt et al. 2009 (Schmidt et al., 2009), with some modifications. First, 6–8 mg of liver (from 2–4 fish) was cut into small pieces and transferred to 1.4 ml of PBS. Then, cross-linking was performed by adding formaldehyde to a final concentration of 1% (v/v) and rocking at room temperature for 8 min. Cross-linking reaction was quenched by adding glycine to a final concentration of 125 mM. To prevent protein degradation, 20 mM sodium butyrate, 1 mM PMSF and 1×cOmplete^TM^ EDTA-free Protease Inhibitor Cocktail (Roche) were added throughout the above procedures. After washing with PBS twice, tissue was dounced first with the loose pestle 20 times and later with the tight pestle 50 times (DWK Life Sciences Inc.) in ice-cold lysis buffer 1 (50 mM HEPES-KOH, pH = 7.6; 140 mM NaCl; 1 mM EDTA; 10% glycerol; 0.5% NP-40; 0.25% Triton-X100; 20 mM sodium butyrate; 1 mM PMSF; 1×cOmplete). After incubation on ice for 10 min, lysate was filtered with pluriStrainer Mini 40 µm (pluriSelect) and washed with the lysis buffer 1 once. The pellet was dissociated with the lysis buffer 2 (10 mM Tris-HCl, pH = 8.0; 200 mM NaCl; 1 mM EDTA; 0.5 mM EGTA; 20 mM sodium butyrate; 1 mM PMSF; 1×cOmplete), rocked at 4℃ for 5 min, and centrifuged. Pellets were dissociated with lysis buffer 3 (50 mM Tris-HCl, pH = 8.0; 10 mM EDTA; 1% SDS; 20 mM sodium butyrate; 1 mM PMSF; 1×cOmplete) and sonicated to sizes ranging from 100–500 bp using Covaris S220 with the following parameters: Peak Power, 105; Duty Factor, 4.0; cycles per burst, 200; duration, 12.5 min.

As a positive control for ChIP experiments, zebrafish chromatin lysate derived from BRF41 fibroblasts was added so that the ratio of medaka and zebrafish DNA was approximately 10:1. Chromatin lysates were diluted with RIPA buffer (10 mM Tris-HCl, pH = 8.0; 140 mM NaCl; 1 mM EDTA; 1 mM EGTA; 1% Triton X-100; 0.1% SDS; 0.1% Na-Deoxycholate; 20 mM sodium butyrate; 1 mM PMSF; 1×cOmplete protease inhibitor) and rocked with the antibody/protein A Dynabeads complex overnight at 4℃. After washing three times with RIPA buffer and once with TE buffer (10 mM Tris-HCl, pH = 8.0; 10 mM EDTA, pH = 8.0), chromatin was de-crosslinked by incubating in lysis buffer 3 at 65℃ overnight. After removing beads, eluted samples were treated with RNase A for two hours at 37℃, followed by proteinase K for two hours at 55℃. DNA was purified by phenol/chloroform extraction and ethanol precipitation. Input DNA was simultaneously treated from the elution step. ChIP-seq libraries were constructed using KAPA Hyper Prep Kit (KAPA Biosystems). Antibodies used in this ChIP experiment are as follows; H3K27ac, ab4729 (abcam); H3K27me3, C15410069 (diagenode); H3K9me3, ab8898 (abcam). Libraries were sequenced using the Illumina HiSeq 1500 platform.

### Immunofluorescence western blotting

Dissected livers were weighed and homogenized in 1×Laemmli sample buffer to the concentration of 1 mg tissue/50 µl buffer. Lysates were separated by 10% polyacrylamide gels and transferred onto PVDF membranes (Immobilon-FL, Millipore). After blocking with 5% skim milk/TBS for 1h at room temperature, membranes were incubated with primary antibodies (rabbit anti-H3K27ac antibody (abcam ab4729), rabbit anti-H3K27me3 antibody (diagenode C15410069) or rabbit anti-H3K9me3 antibody (abcam ab8898) combined with mouse anti-H3 antibody (active motif 39763), 1:2000 dilution each) overnight at 4℃. After several washes, membranes were incubated with secondary antibodies (IRDye 800CW Donkey anti-rabbit IgG (LI-COR) and IRDye 680RD Donkey anti-mouse IgG (LI-COR), 1:10000 dilution) for one hour at room temperature. Visualization of protein bands and quantification of signal intensities were performed using ODYSSEY CLx (LI-COR). The quantity of each histone modification was calculated by dividing the signal intensity of histone modification by the signal intensity of histone H3.

### RNA-seq data processing

Sequenced reads were preprocessed to remove low-quality reads/bases using Trimmomatic v0.33 (Bolger et al., 2014) with the following parameters: SLIDINGWINDOW:4:15, LEADING:20, TRAILING:20, MINLEN:20. Trimmed reads were mapped to the Hd-rR medaka reference genome (Ensembl ASM223467v1.95 (Ichikawa et al., 2017)) by STAR (Dobin et al., 2013). Reads with mapping quality (MAPQ) larger than or equal to 20 were used for further analyses. Differentially expressed genes were obtained by DESeq2 (Love et al., 2014), and gene ontology analyses of differentially expressed genes were performed using PANTHER v17.0 (Thomas et al., 2022), putting all expressed genes in medaka hepatocytes as background.

### ATAC-seq data processing

Sequenced reads were preprocessed to remove low-quality reads/bases using Trimmomatic v0.33 (Bolger et al., 2014) with the following parameters: SLIDINGWINDOW:4:15, LEADING:20, TRAILING:20, MINLEN:20. Trimmed reads were mapped to the Hd-rR medaka reference genome (Ensembl ASM223467v1.95 (Ichikawa et al., 2017)) by BWA (Li and Durbin, 2009). Reads with mapping quality (MAPQ) larger than or equal to 20 were used for further analyses. Peak calling was performed for each data by MACS2 v2.1.0 (Zhang et al., 2008) with the following options: --nomodel --extsize 200 --shift -100 -g 600000000 -q 0.01 -B --SPMR. Peaks common to at least six of the 28 samples (30,484 in total) were used for further analyses. The number of reads in each peak was counted by annotatePeaks command of HOMER (Heinz et al., 2010), and differentially accessible peaks were obtained by DESeq2, based on the read counts inside peaks. Motif analyses of differentially accessible peaks were performed using HOMER. Annotation of the peaks to nearby genes was performed by the following definition: Firstly, ATAC-seq peaks were annotated with the genes in which TSS is located within 2 kb of either end of the ATAC-seq peak. If no TSS was found within 2 kb of either end of the ATAC-seq peak, then we annotated the ATAC-seq peaks with the overlapping genes (if any). ATAC-seq peaks can be associated with multiple genes when several TSSs or genes are closely located (e.g., bidirectional promoters). Peaks that are located inside 2 kb on either side of the TSS are categorized as promoters, and the others as enhancers. Gene ontology analyses of genes near differentially accessible peaks were performed using PANTHER v17.0 (Thomas et al., 2022), putting all genes annotated to at least one of the ATAC-seq peaks as background (13,622 genes in total).

### ChIP-seq data processing

Sequenced reads were preprocessed to remove low-quality reads/bases using Trimmomatic v0.33 (Bolger et al., 2014) with the following parameters: SLIDINGWINDOW:4:15, LEADING:20, TRAILING:20, MINLEN:20. Trimmed reads were aligned to the concatenated Hd-rR medaka (Ensembl ASM223467v1.95 (Ichikawa et al., 2017)) and zebrafish (Ensembl GRCz11.95) genomes by BWA (Li and Durbin, 2009). Reads mapped to the medaka genome with mapping quality (MAPQ) larger than or equal to 20 were used for further analyses. For H3K27ac, peak calling was performed for each dataset by MACS2 v2.1.0 (Zhang et al., 2008) with the following options: -- broad -g 600000000 -q 0.01 -B --SPMR. Then peaks were merged for all the 12 datasets, and 35,722 peaks were obtained in total. Differential analyses, gene annotation, and gene ontology analyses were performed in the same methods as ATAC-seq. For H3K27me3 and H3K9me3, differentially enriched regions were obtained by dividing the medaka genome into 4-kb bins, counting the number of reads at each bin, and performing DESeq2.

### Normalization and visualization of bedgraphs

Counts per million (CPM)-normalized bedgraphs were generated by MACS2 (Zhang et al., 2008) with the following options. ATAC-seq, -g 600000000 -q 0.01 -B -- SPMR --nomodel --extsize 200 --shift -100; H3K27ac ChIP-seq, --broad -g 600000000 - q 0.01 -B --SPMR; H3K27me3 and H3K9me3 ChIP-seq, --broad -g 600000000 -q 0.01 - B --SPMR --nomodel --extsize 190. Owing to the variation of peak height in the genome browser view, possibly due to the difference in signal/noise ratio among samples, we further performed normalization of ATAC-seq and ChIP-seq track data (Figure 5–figure supplement 2). First, we determined the scaling factors for visualization of ChIP-seq/ATAC-seq data by linear regression procedure as follows. Here, we only used regions with stable signals (i.e., read count). Started with all 2 kb regions around TSS (for ATAC-seq, and K27ac marks) or peak regions consistent across the libraries (for K27me3 and K9me3 marks), we excluded a region if the average signal was within the top 100 or below the global average (average across all regions and libraries). We also excluded a region if the variation of signals was within the top 10%. Then, the signals of the remaining regions for each library were regressed against the averages across libraries. The resulting coefficient (slope) was identified as the scaling factor for each library. Finally, we divided the track data by the calculated scaling factors.

## Author contributions

Y.I., S.M., and H.T. planned and supervised this research. Y.I. and Y.K. established fish-rearing conditions. Y.K. performed histological analyses of liver tissues. Y.K. created the *tdo2*-GFP transgenic medaka. Y.I. and Y.K. performed RNA-seq library preparation and sequencing. Y.K. performed ATAC-seq library preparation and sequencing. Y.I. and T.W. performed ChIP-seq library preparation and sequencing. Y.I. performed western blotting. Y.I., Y.S., and Y.K. performed data analyses of RNA-seq, ATAC-seq, and ChIP-seq data. Y.S. prepared normalized track views of ATAC-seq and ChIP-seq data. Y.I. and H.T. wrote the manuscript.

## Acknowledgments

We thank Prof. K. Kamimura, Dr. R. Goto, and Prof. S. Terai for the help in establishing fish rearing conditions, Dr. Y. Kimura and Prof. S. Higashijima for kindly providing us with plasmids for CRISPR-Cas9-mediated gene knockin, and Mrs. M. Sakamoto for fish care. We acknowledge Tokai Electron Microscopy for performing electron micrographs of liver tissue. Flow cytometry was performed in the IMSUT FACS Core laboratory. We acknowledge the IMSUT FACS Core laboratory for assistance with flow cytometry analysis. (FACS). This work was supported by AMED CREST, JST (Grant No: JP23gm1110007).

## Data access

Sequencing data generated in this study has been submitted to the DDBJ BioProject database under accession number PRJDB15283.

---

**Figure 2–figure supplement 1:**
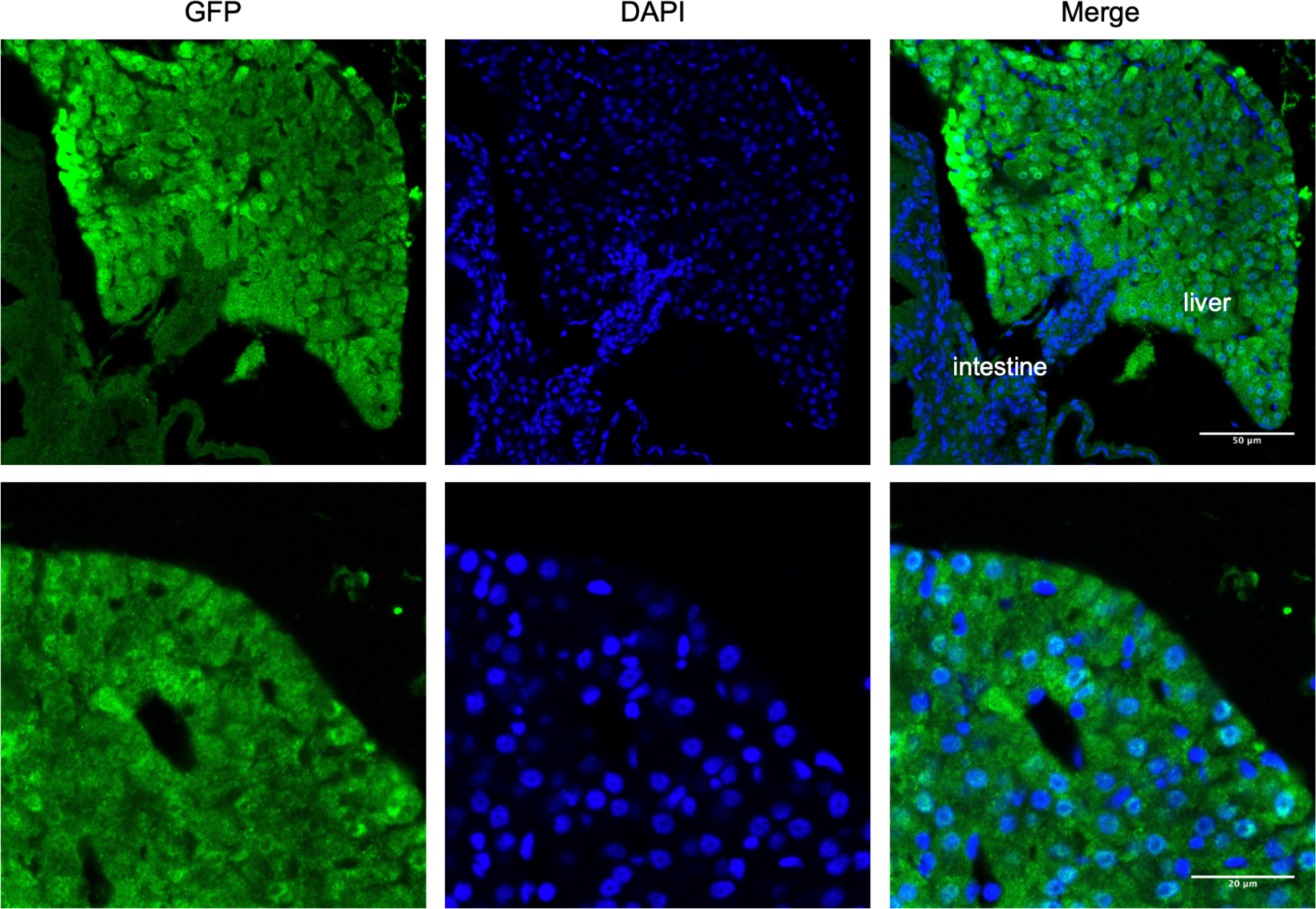
Confirmation of hepatocyte-specific expression of GFP in *tdo2*:GFP transgenic medaka. Immunofluorescent images of a liver section stained with GFP antibody are shown. Note that GFP signal was detected in hepatocytes, but not in red blood cells, endothelial cells in the liver, or cells in the intestine. Scale bar: 50 µm (upper), 20 µm (lower).

**Figure 2–figure supplement 2:**
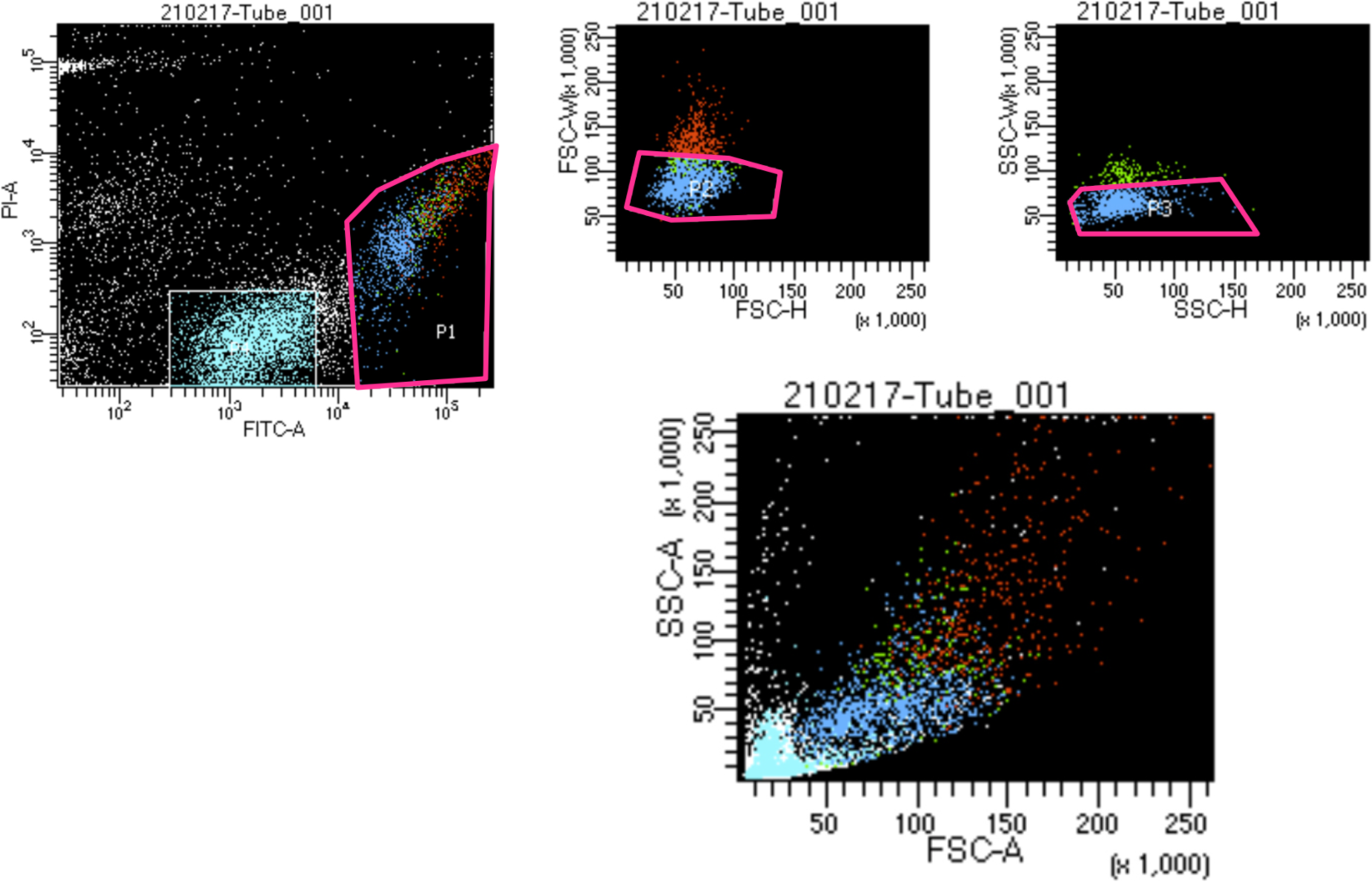
Sorting of GFP-positive hepatocytes from livers of *tdo2*:GFP medaka. FACS-sorting strategy of GFP-positive hepatocytes from liver cell suspensions of *tdo2*:GFP transgenic medaka. GFP-positive, propidium-iodide (PI)-negative, singlet cells were sorted.

**Figure 2–figure supplement 3:**
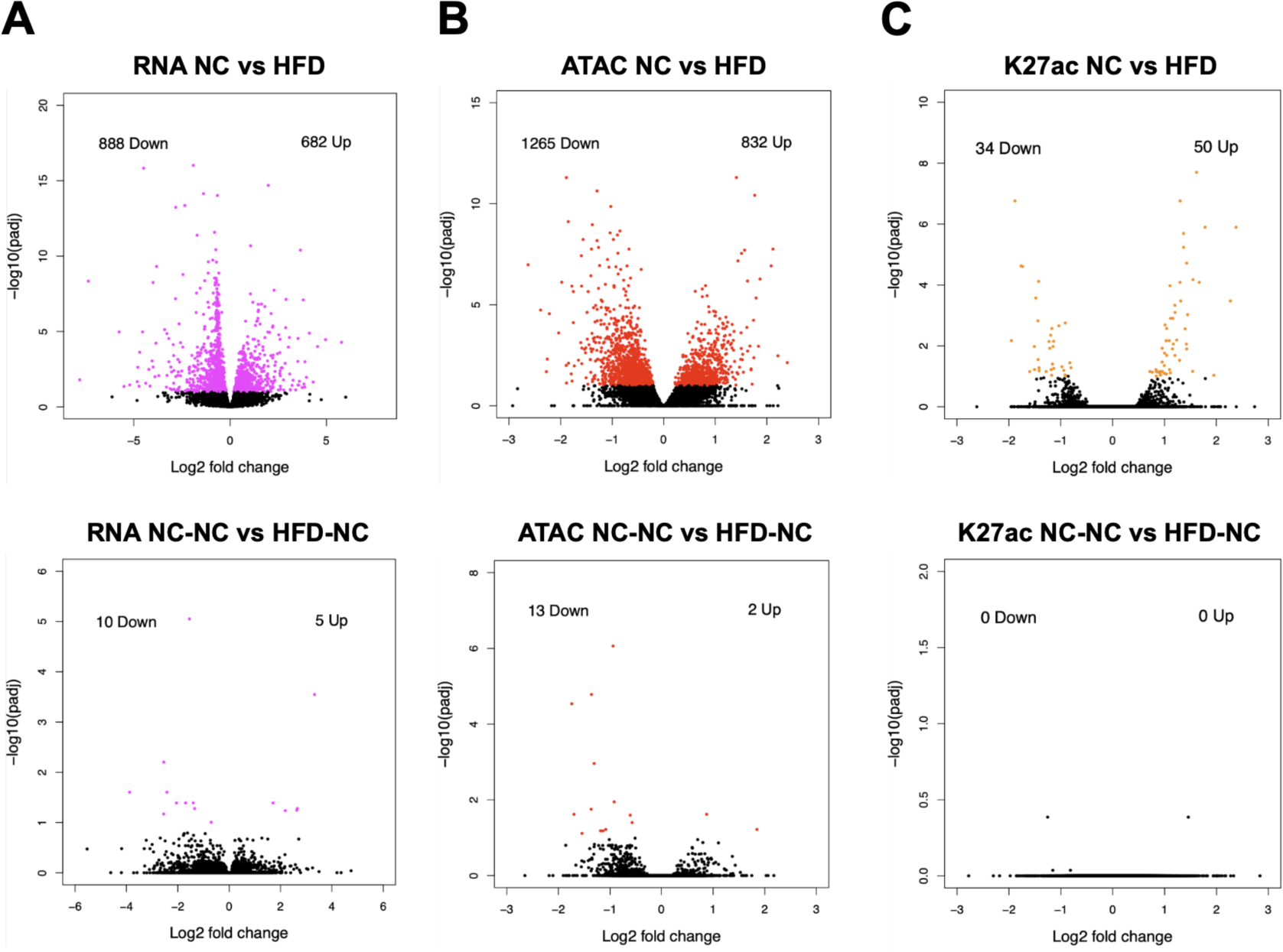
Differentially expressed genes/peaks at FDR < 0.1, corrected for multiple testing. Volcano plots showing **(A)** differentially expressed genes by RNA-seq, **(B)** differentially accessible peaks by ATAC-seq, and **(C)** differentially H3K27ac-enriched peaks by ChIP-seq. X-axis: log2 fold change of normalized read counts within genes/peaks (upper, HFD/NC; lower, HFD-NC/NC-NC), y-axis: –log10(FDR).

**Figure 3–figure supplement 1:**
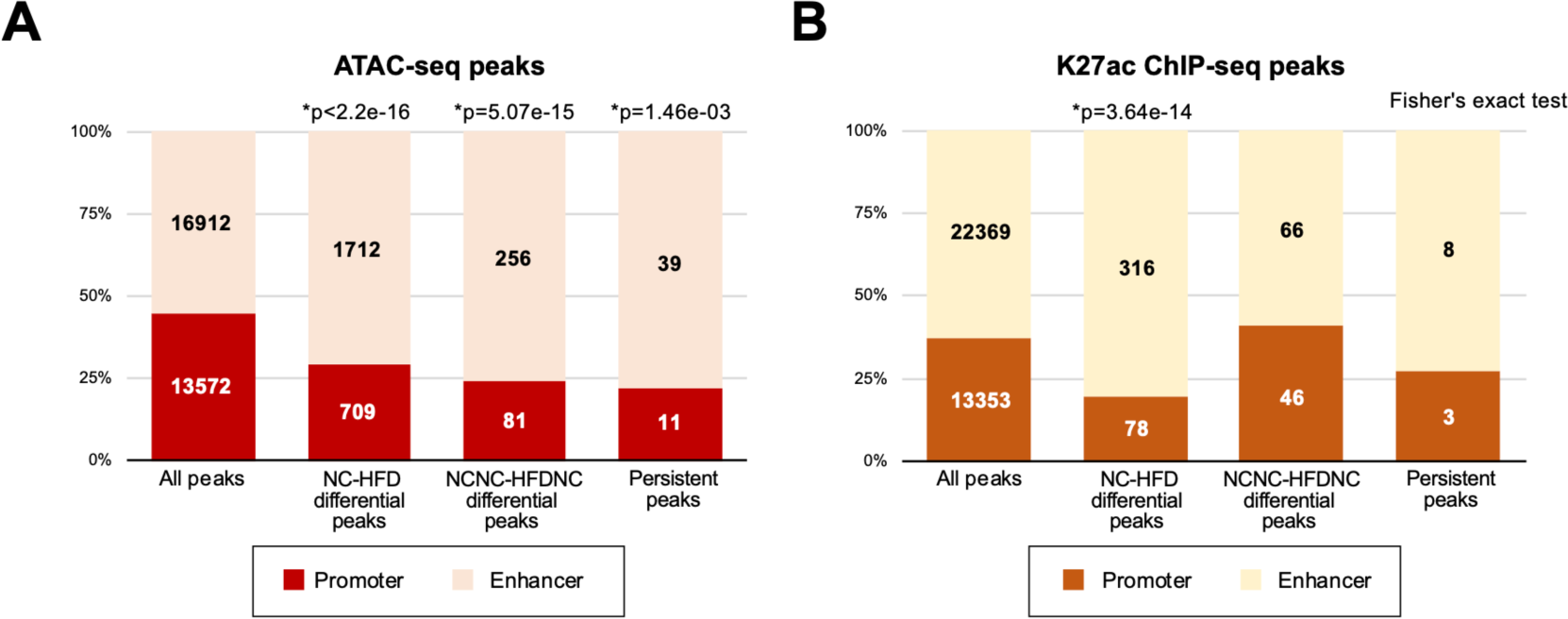
Categorization of peaks into promoters and enhancers. **(A)** ATAC-seq peaks (30,484 peaks in total) were categorized as promoters if peaks were located inside 2 kb on either side of the transcription start sites (TSS), while others were categorized as enhancers. **(B)** H3K27ac ChIP-seq peaks (35,722 peaks in total) were categorized as promoters if peaks were located inside 2 kb on either side of the TSS, while others were categorized as enhancers. Enrichment of peaks with differential signal at enhancers was tested by Fisher’s exact test.

**Figure 4–figure supplement 1:**
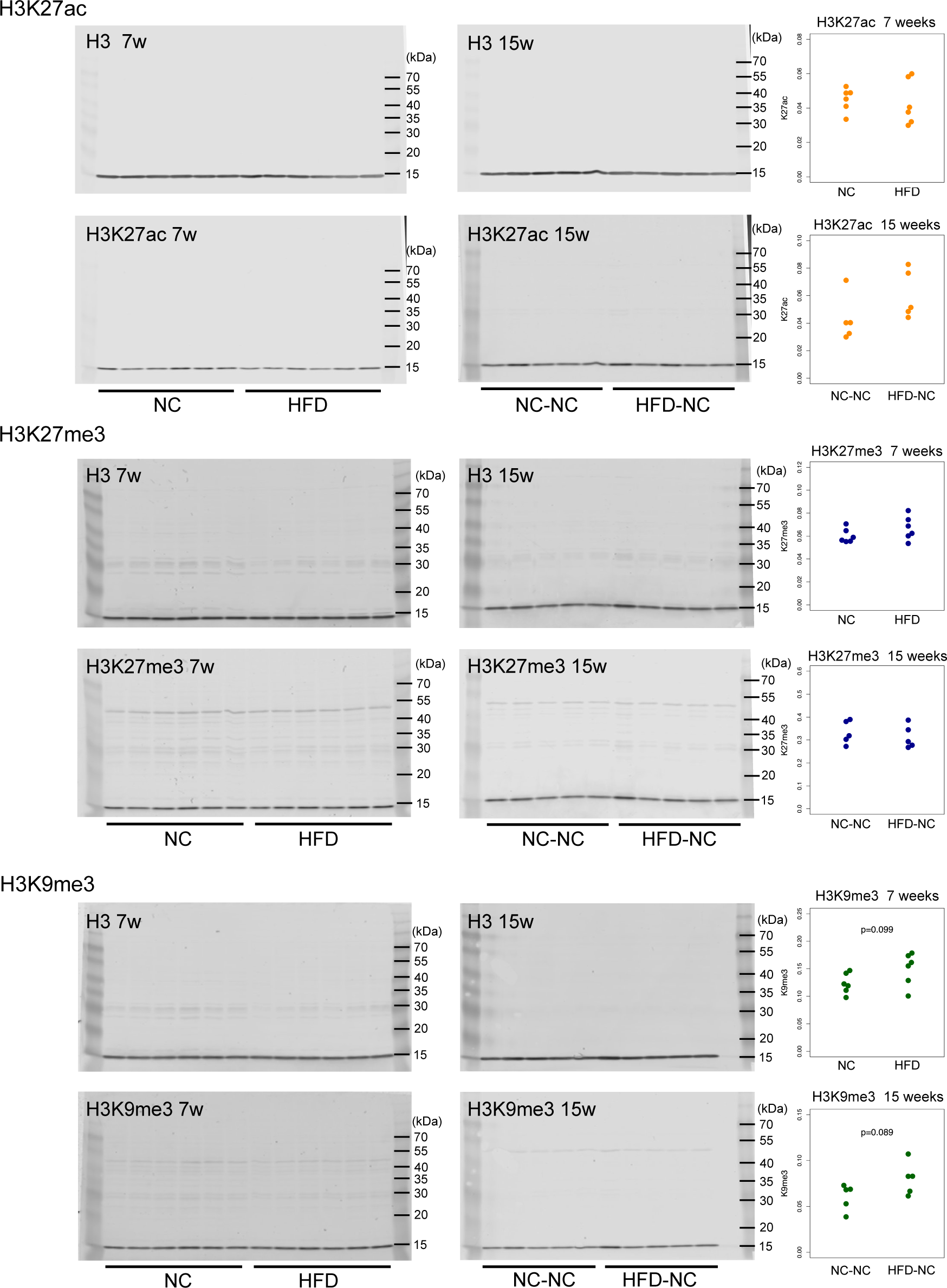
Western blotting of H3K27ac, H3K27me3, and H3K9me3 histone modifications in livers. Left: Raw images of western blotting of each histone modification and histone H3. One lane indicates liver lysate of a single fish, concentration of which was adjusted by liver weight. Right: plots of signal intensities of each histone modification, normalized by signal intensities of histone H3. Note that there is little difference in the global levels of each histone modification in medaka liver between dietary conditions.

**Figure 4–figure supplement 2:**
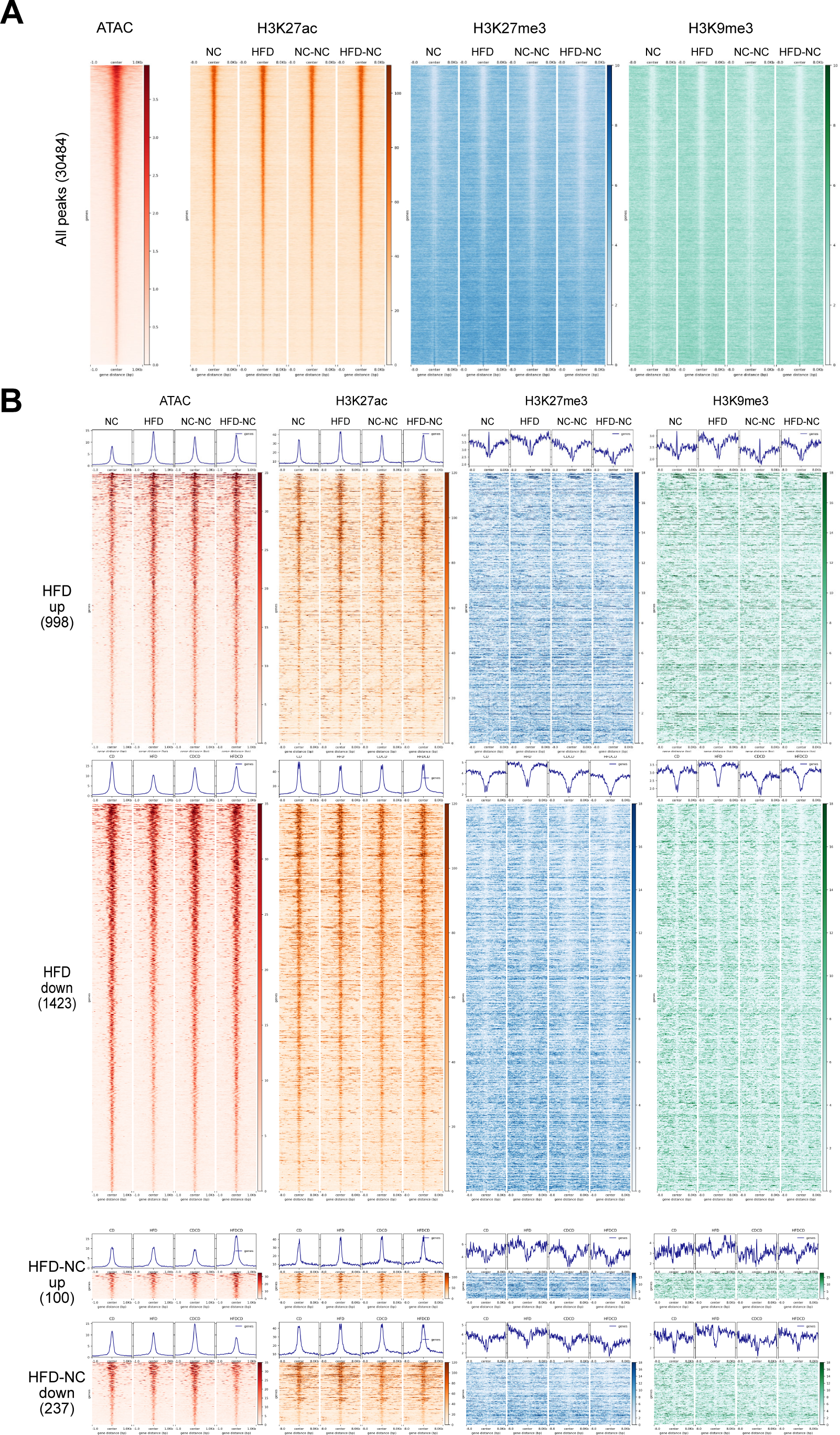
Distribution pattern of histone modifications around ATAC-seq peaks. **(A)** Distribution pattern of H3K27ac (orange), H3K27me3 (blue), and H3K9me3 (green) around all 30,484 ATAC-seq peaks, sorted by ATAC-seq signal intensity. For ATAC-seq, signal intensities within a 1 kb window from the centers of ATAC-seq peaks are displayed. For histone modifications, signal intensities within an 8 kb window from the centers of ATAC-seq peaks are displayed. Note the positive correlation of ATAC-seq signal with H3K27ac, and the negative correlation with H3K27me3 and H3K9me3 signal. **(B)** Distribution patterns of each histone modification around differentially accessible peaks. For peaks differentially accessible between HFD and NC fish, a moderate positive correlation was observed for H3K27ac. However, the correlation was not observed for differentially accessible peaks between HFD-NC and NC-NC fish. For H3K27me3 and H3K9me3, little changes were seen for differentially accessible peaks at any time point.

**Figure 4–figure supplement 3:**
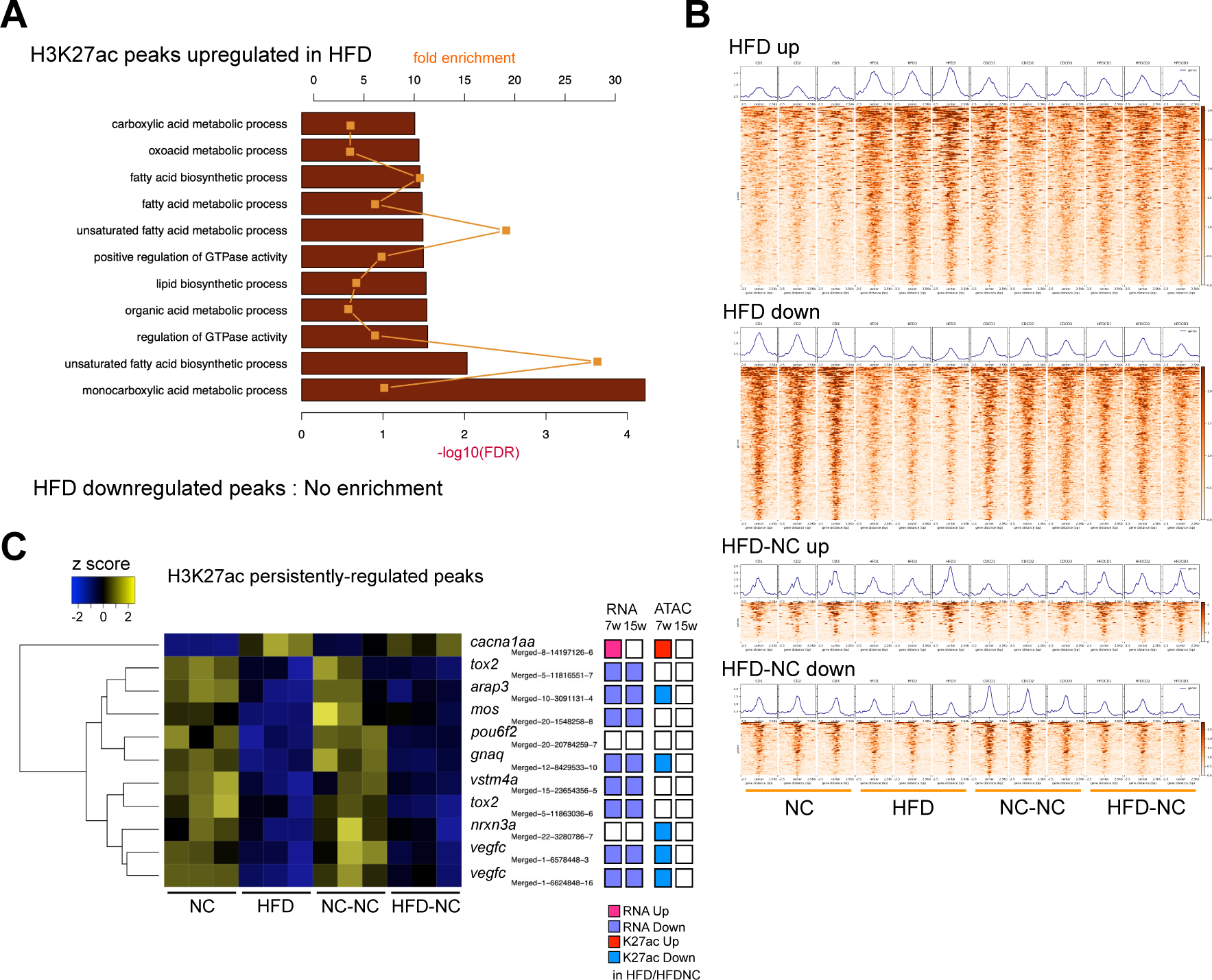
Differentially enriched H3K27ac peaks by HFD feeding. **(A)** Gene ontology analyses of genes close to H3K27ac peaks with different enrichment between HFD and NC group fish, inferred by PANTHER v17.0. **(B)** Heatmaps of H3K27ac peaks differentially enriched at seven and 15 weeks of age. Signal intensities in ±2.5 kb regions from the centers of H3K27ac peaks are displayed. **(C)** A heatmap of persistent H3K27ac ChIP-seq peaks after NC (11 peaks in total). Log2-transformed, and Z-transformed, DESeq2 normalized read counts at each peak were displayed. DESeq2 results of RNA-seq and ATAC-seq of nearby genes/peaks were displayed on the right.

**Figure 4–figure supplement 4:**
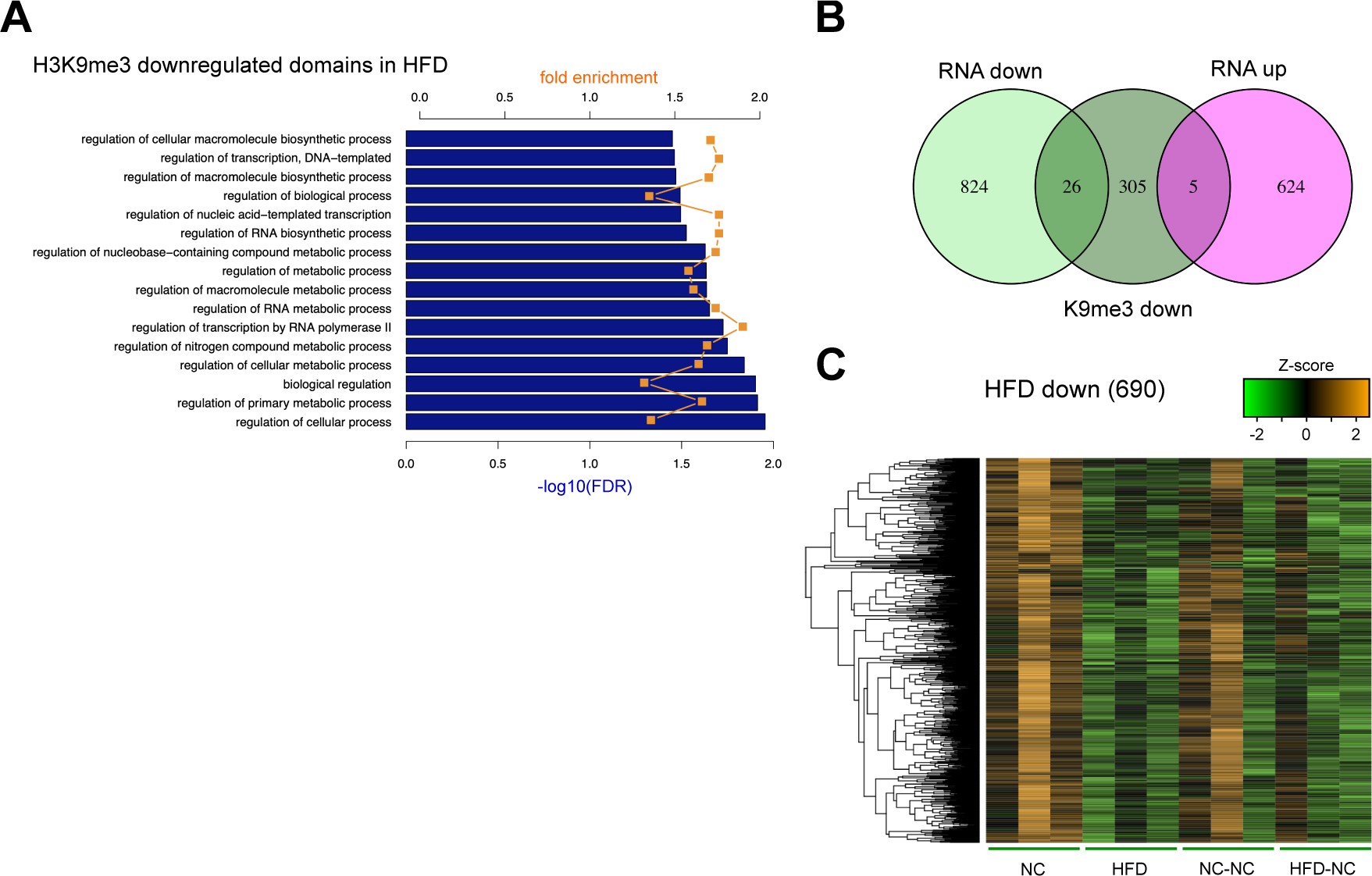
H3K9me3 domains downregulated by HFD feeding. **(A)** Gene ontology analyses of genes close to H3K9me3 bins differentially enriched between HFD and NC group fish. **(B)** Venn diagram of genes close to H3K9me3-downregulated bins and genes differentially expressed after HFD feeding. **(C)** A heatmap of H3K9me3 levels for the downregulated bins by HFD. Log2-transformed, and Z-transformed, DESeq2 normalized read counts of H3K9me3 ChIP-seq at each bin are displayed.

**Figure 5–figure supplement 1:**
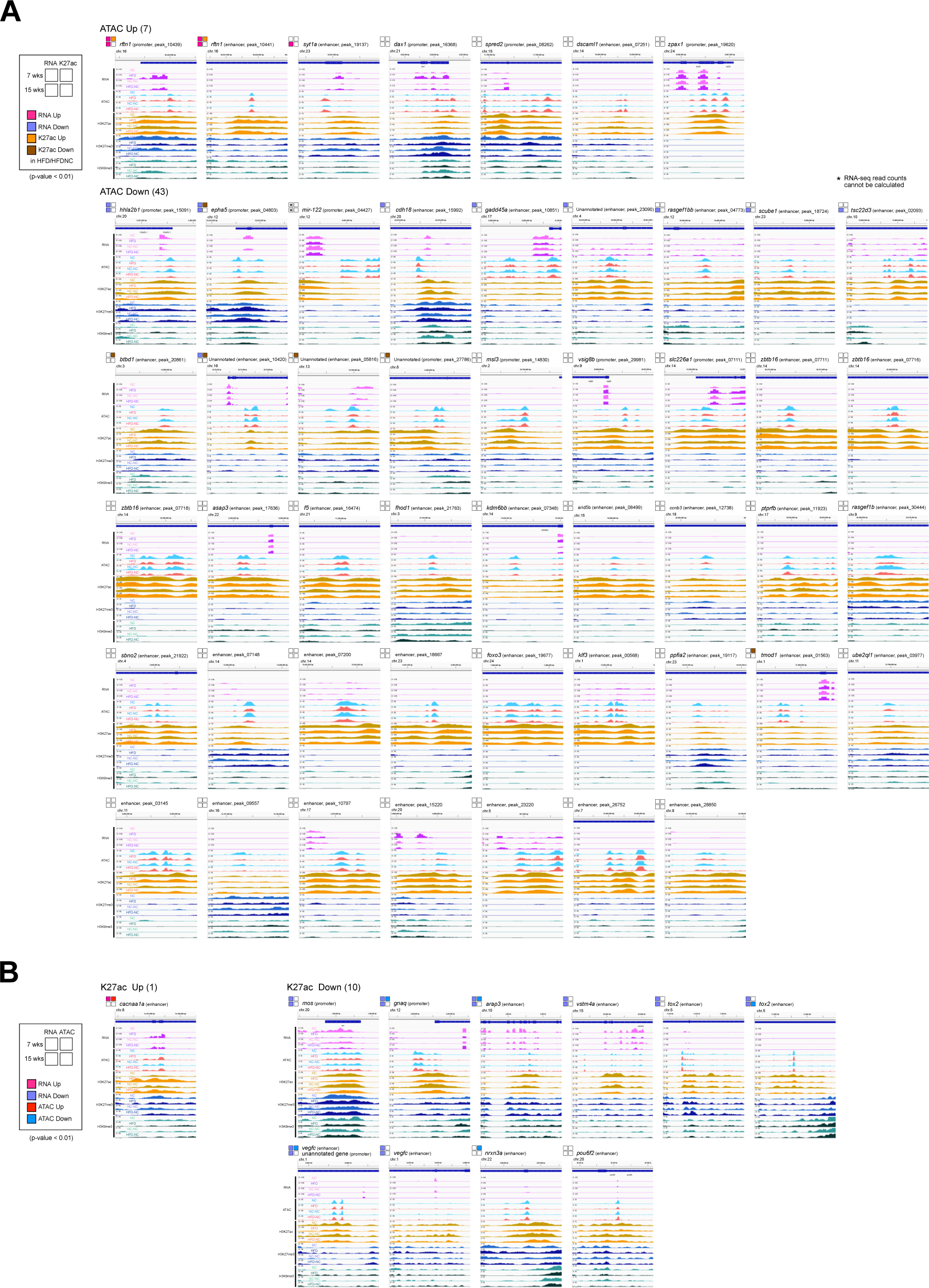
Track views of persistent peaks. **(A)** Track views of the 50 persistent ATAC-seq peaks. DESeq2 results of RNA-seq and H3K27ac ChIP-seq of nearby genes/peaks are displayed on the upper left. **(B)** Track views of the 11 persistent H3K27ac ChIP-seq peaks. DESeq2 results of RNA-seq and ATAC-seq of nearby genes/peaks are displayed on the upper left.

**Figure 5–figure supplement 2:**
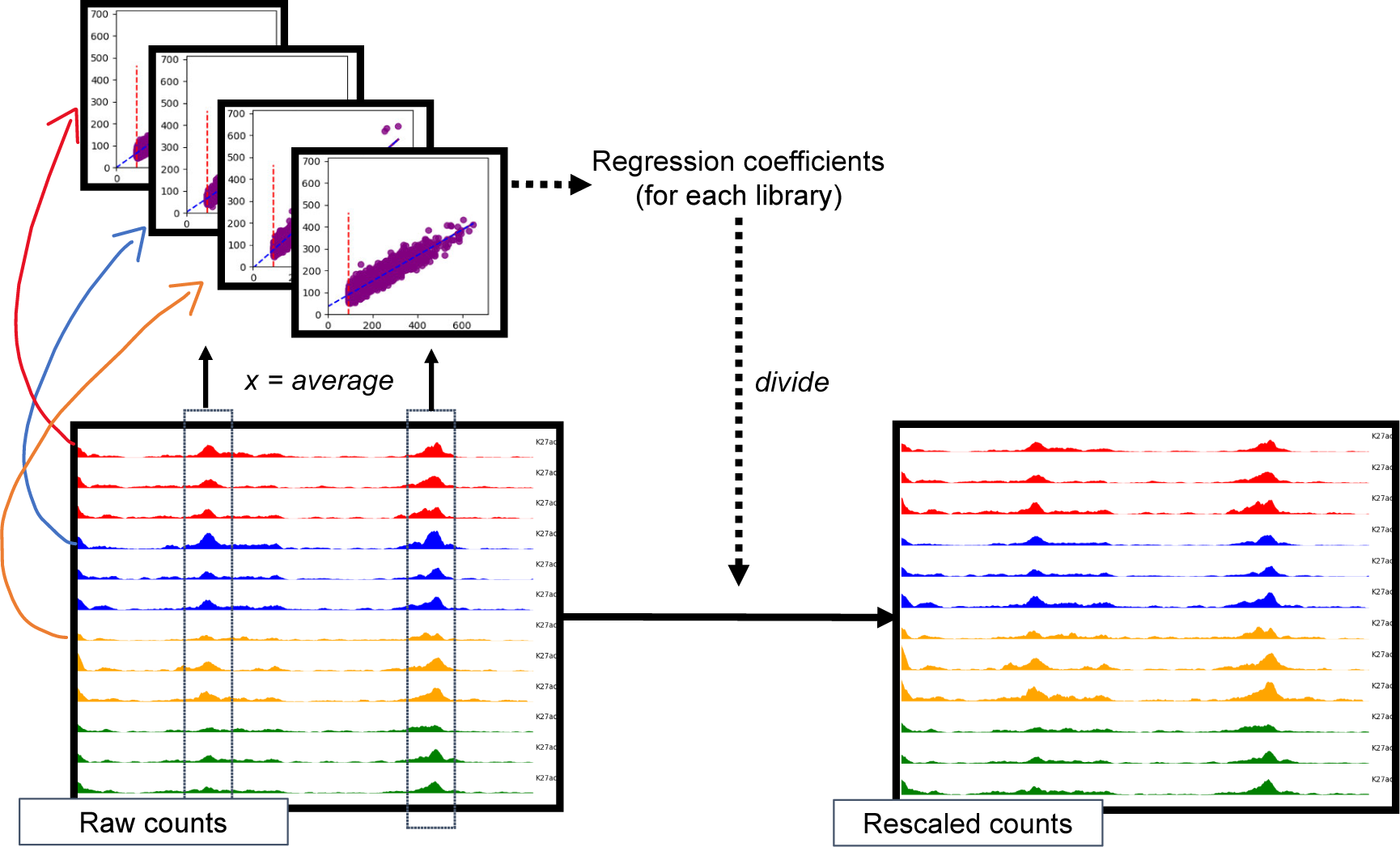
Rescaling procedure of read counts for visualization of ChIP-seq and ATAC-seq. The top left panels describe the regression process. Each point in these panels represents read count data from a single region. For each library, the signals on the selected regions (y-axis) were regressed on the averages of all libraries (x-axis). In this process, the regions whose average signal was within the top 100 or below the global average (average across all regions and libraries) were excluded (red dotted lines). Then, the raw counts (left bottom panel) were divided by the corresponding regression coefficients (blue dotted lines) to obtain the rescaled counts (the right bottom panel).

**Supplementary File 1: List of genes differentially-expressed / differentially-accessible / differentially-H3K27ac-enriched by diet.**

